# Specialized medial prefrontal-amygdala coordination in other-regarding decision preference

**DOI:** 10.1101/640292

**Authors:** Olga Dal Monte, Cheng-Chi J. Chu, Nicholas A. Fagan, Steve W. C. Chang

## Abstract

Social behaviors recruit multiple cognitive processes requiring coordinated interactions among brain regions. Oscillatory coupling provides one mechanism for cortical and subcortical neurons to synchronize their activity. However, it remains unknown how neurons from different nodes in the social brain network interact when making social decisions. We investigated neuronal coupling between the rostral anterior cingulate gyrus of the medial prefrontal cortex and the basolateral amygdala while monkeys expressed context-dependent positive other-regarding preference (ORP) or negative ORP impacting the reward of another monkey. We found an enhanced synchronization between the two nodes for positive ORP, but a suppressed synchronization for negative ORP. These interactions occurred in dedicated frequency channels depending on the area contributing spikes, exhibited a specific directionality of information flow associated with expressing positive ORP, and could be used to decode social decisions. These findings support that specialized coordination in the medial prefrontal-amygdala network underlies social decision preference.

---

Altruistic behaviors and mutually-beneficial social exchanges facilitate social cohesion among members of a group and help attain collective rewards. While selfish behaviors can be detrimental to these causes, they may be strategically necessary to secure limited resources or achieve a certain social status. The cognitive operations central to making such social decisions are theorized to recruit a wide array of brain regions that are sensitive to primary and more abstract rewards, and span both cortical and subcortical areas with divergent functional specifications^1–5^.

Recent single-neuron studies using social decision-making paradigms involving pairs of monkeys have begun to characterize neuronal correlates of social decision variables concerning conspecific animals in several brain regions. These regions include the anterior cingulate cortex (ACC)^6, 7^, the dorsomedial prefrontal cortex^8^, the basolateral amygdala (BLA)^9–11^, the orbitofrontal cortex (OFC)^6, 12^, the striatum^13^, as well as the lateral prefrontal cortex^14, 15^. Of these, the gyrus of the rostral ACC (ACCg) of the medial prefrontal cortex is thought to be particularly specialized in signaling rewarding and motivational information about social partners in both humans and monkeys^1, 16^. Specifically, in a task where monkeys can express their other-regarding preferences (ORP) by choosing to deliver juice rewards to a conspecific monkey over discarding the rewards, some ACCg cells exclusively encode conspecific’s rewards while other cells encode one’s own reward and conspecific’s reward in an indistinguishable manner^6^. By contrast, neurons in the OFC or in the sulcus of the ACC in the same paradigm predominantly signal self-referenced decision variables by modulating firing rates only in relation to one’s received or foregone rewards^6^. These findings lend support for the role of rostral ACCg in computing other-referenced decisions ^16^. On the other hand, BLA neurons, in the same behavioral task, exhibit systematically correlated firing rates for encoding monkeys’ choices that result in juice rewards allocated to either themselves or the conspecific monkey^9^, suggesting that subcortical neurons in BLA utilize a shared metric for computing decision variables across self and other. These general neural characteristics in relation to social decision variables have also later been observed in ACCg and BLA neurons in the human brain in an intracranial study^17^. While brain regions like ACCg and BLA are implicated in social decision-making, it is likely that the systematic synchronization across these and similar brain regions is what truly underlies such decisions.

Specialized coherence signature across specific nodes in the social brain network likely plays a key role in social cognition. Whole-brain functional neuroimaging studies in humans have indicated the potential importance of correlated hemodynamic fluctuations across different brain regions in regulating complex social cognition^18, 19^. In prairie voles, frequency-specific coupling between medial prefrontal cortex and nucleus accumbens is shown to mediate social bonding^20^. Moreover, ACC neurons in the medial prefrontal cortex network that directly project to BLA are found to be necessary for observational fear learning and social preference formation in mice^21^. In turn, dysregulated subcortical-medial prefrontal synchrony could result in abnormal social behaviors^22^. However, while there is growing evidence for the importance of interactive coordination, neuronal mechanisms underlying interareal synchrony associated with complex social behaviors, such as those related to positive or negative ORP, remain elusive.

Reciprocally and densely innervating anatomical projections between ACCg and BLA permit the two nodes to efficiently communicate with one another for processing social and affective information^23, 24^. However, whether and how ACCg and BLA coordinate their activity in relation to social decision-making remain unknown. If coordinated interactions between ACCg and BLA were involved in the expression of either positive or negative ORP concerning the welfare of others, one might expect distinctive coordination patterns to exist for two different types of expressed ORPs. Such coordinated interaction may be mediated by a dedicated frequency channel with a specific information flow between ACCg and BLA associated with expressing social decision preferences. To directly test this hypothesis, we investigated how single-neuron spiking and local field potential (LFP) activity between ACCg and BLA are dynamically coordinated as monkeys expressed positive ORP or negative ORP toward a conspecific monkey. We used spike-field coherence as our primary measure as it quantifies how spiking activity from one brain region is synchronized to oscillatory LFP activity from another brain region in discrete time and frequency windows, allowing inspections of synchronous coordination of neural activity across brain areas^25, 26^.

We found that synchrony between spiking and LFP oscillations in the two nodes differentiated monkeys’ positive ORP in one context (via enhanced spike-field coherence) from negative ORP in another context (via suppressed spike-field coherence). Moreover, these synchrony patterns were specific to select frequency bands and time windows, and support a directional relationship of information transfer between the two nodes. Taken together, our findings demonstrate that unique rhythmic coordination of neuronal activity in the primate medial prefrontal-amygdala network contributes to context-specific social decision-making.

## Results

### Monkeys exhibit positive ORP and negative ORP in distinct contexts

Pairs of rhesus macaques (an actor and a recipient) participated in the social reward allocation task (**Fig. 1a-b**; Online Methods). In one decision context (*Other*/*Bottle* context) where actor monkeys never received juice rewards, actors were free to choose between donating a juice drop to a recipient (*Other*) and to a juice collection bottle (*Bottle*). In the other decision context (*Self*/*Both* context) where actors always received juice rewards, actors were free to choose between delivering juice rewards to themselves (*Self*) and to both themselves and the other monkey (*Both*). There were three magnitudes of juice reward offered (Online Methods), and actors were informed of the value at stake on each trial. This task therefore measures actor’s social decision preference without self-reward confound in choosing one option over the other in two separate contexts.

**Fig. 1.**
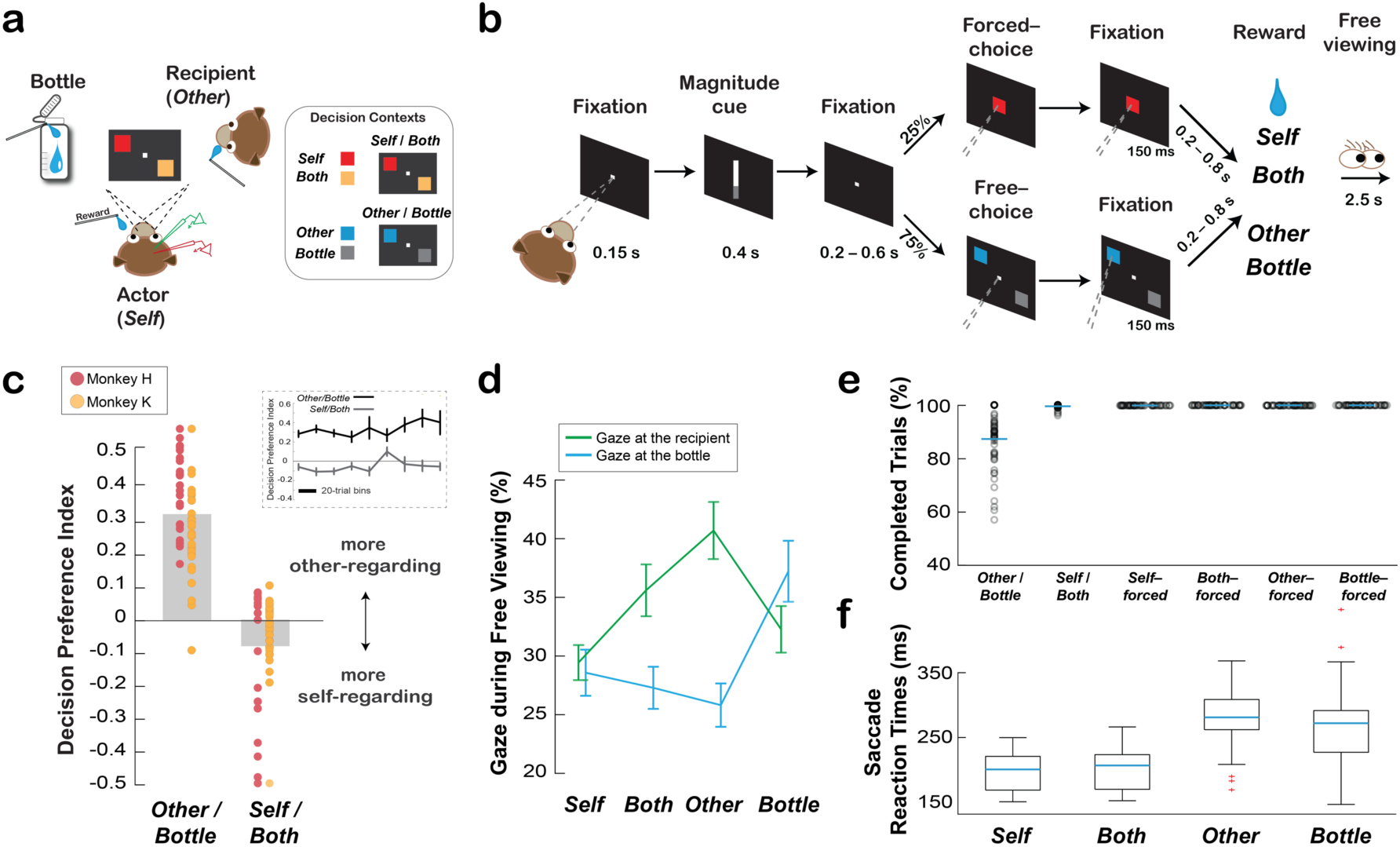
Social reward allocation task and the behaviors associated with social decision preference. (**a**) Experimental setting involving an actor monkey, a recipient monkey, and an operating juice collection bottle. The inset shows example stimulus-reward outcome mappings for the two distinct contexts for rewarding the actor (*Self*) or both the actor and the recipient (*Both*) (*Self*/*Both* context), and for rewarding the recipient (*Other*) or the bottle (*Bottle*) (*Other*/*Bottle* context). (**b**) Task sequence for the social reward allocation task (Online Methods). (**c**) Monkeys exhibited context-dependent positive and negative ORPs. Decision preferences are expressed as averaged contrast ratios for the two decision contexts. Data points overlaid on top show the biases for all individual sessions for each subject. The inset shows the preferences over time for each context. (**d**) Social gaze patterns reflected decisions to deliver juice rewards to the recipient or the bottle as a function of different decisions. Shown are the mean (± s.e.m.) proportions of gaze to the recipient or to the bottle during the free viewing period for each reward outcome. (**e**) Average proportions of completed free-choice trials for *Other*/*Bottle* and *Self*/*Both* contexts and completed forced-choice trials for choosing *Self*, *Both*, *Other*, or *Bottle*. Data points show individual sessions. (f) Saccade reaction times (mean ± s.e.m.) for choosing *Self*, *Both*, *Other*, or *Bottle*.

Actors completed 313 ± 109 (mean ± s.d.) trials per session across 57 sessions (monkey H: 374 ± 110 per session, 31 sessions; monkey K: 240 ± 43 per session, 26 sessions). Consistent with previous findings using this behavioral design^6, 9, 27, 28^, actors preferred to choose *Other* over *Bottle*, exhibiting a positive ORP (preference index, mean ± s.e.m.: 0.32 ± 0.02, p < 0.0001, Wilcoxon sign rank) in the *Other*/*Bottle* context, but preferred to choose *Self* over *Both*, displaying a negative ORP in the *Self*/*Both* context (–0.08 ± 0.02, p < 0.001) (**Fig. 1c**). These context-dependent preferences were consistent and stable over time of each session (*Self*/*Both* and *Other*/*Bottle* context: both p > 0.52, linear regression, **Fig. 1c**)^9, 27^, have been observed across several different animals in independent studies^6, 9, 27, 28^, are sensitive to dominance and familiarity between pairs^27^, and are abolished if the recipient monkey is replaced with a juice collection bottle^27^.

Social gaze patterns differed as a function of decision (*Self*, *Both*, *Other*, *Bottle*) (F[3, 455] = 2.86, p = 0.037) and gaze-goal (the recipient or the bottle) (F[1, 455] = 10.66, p = 0.001). Critically, decision type and gaze-goal showed a strong interaction (F[3, 455] = 8.75, p < 0.0001), indicating that social gaze differed across decision types. Across all decision outcomes, actors looked at the recipient (36 ± 1% [mean ± s.e.m.]) at a higher rate than to the bottle (30 ± 1%, p = 0.001, Tukey test). Importantly, after choosing *Other,* actors looked at the recipient (41 ± 2%) more frequently compared to the bottle (26 ± 2%, p < 0.0001). By contrast, actors looked at the bottle more often after choosing *Bottle* (37 ± 3%) than after choosing *Other* (26 ± 2%) (p = 0.002) (**Fig. 1d**). These observations support that actors were acutely aware of the reward outcome differences between the two conditions in which rewards were either allocated to the recipient or the bottle, the two outcomes without a self-reward contingency^6, 9, 27, 28^. These context-dependent social decision preferences of the actor monkeys provide a behavioral framework for examining the coordination between ACCg and BLA in expressing positive and negative ORPs toward a conspecific monkey under different contexts.

As expected during free-choice trials, actors overall completed more *Self/Both* trials (greater than 99% for all reward sizes) compared to *Other/Bottle* trials (87% for all reward sizes) (F[1,341] = 175.12, p < 0.0001) (**Fig. 1e**). However, actors were more motivated to complete *Other*/*Bottle* trials when the reward size at stake for either the recipient or the bottle was larger (small: 83 ± 2%, medium: 87 ± 2%, large: 90 ± 2%; F[2,168] = 4.3, p = 0.02). On forced-choice trials, performance was at ceiling and did not differ across outcomes. Saccade reaction times on free-choice trials differed as a function of decision (*Self* [197 ms ± 27 ms], *Both* [200 ms ± 29 ms], *Other* [278 ms ± 43 ms], *Bottle* [271 ms ± 59 ms]; F[3, 215] = 59, p < 0.0001) (**Fig. 1f**), driven by the differences in reaction times for receiving rewards (*Self* or *Both*) compared to forgoing rewards (*Other* or *Bottle*) (p < 0.0001, Wilcoxon rank sum; *Self* vs. *Both*, *Other* vs. *Bottle*, both p > 0.75; *Self* or *Both* vs. *Other* or *Bottle*, all p < 0.001, Tukey test).

### Coordination of spiking and LFP activity between ACCg and BLA

Exploiting monkeys’ context-dependent positive and negative ORPs, we investigated neural coordination relating spiking and LFP activity associated with the two types of ORPs between rostral ACCg (Brodmann areas 24a, 24b, and 32)^29^ and BLA^29^ (**Fig. 2**). All single units were recorded without any sampling criterion, resulting in 253 ACCg cells and 90 BLA cells. **Figure S1** shows basic characterizations of the single cell activity as well as example cells with outcome selective response profiles. As we have previously characterized single-cell encoding of social decision variables within ACCg and BLA in the identical social reward allocation task^6, 9^, in this study we mainly focused on determining coordination in frequency and time between ACCg and BLA at the level of single cells and populations.

**Fig. 2.**
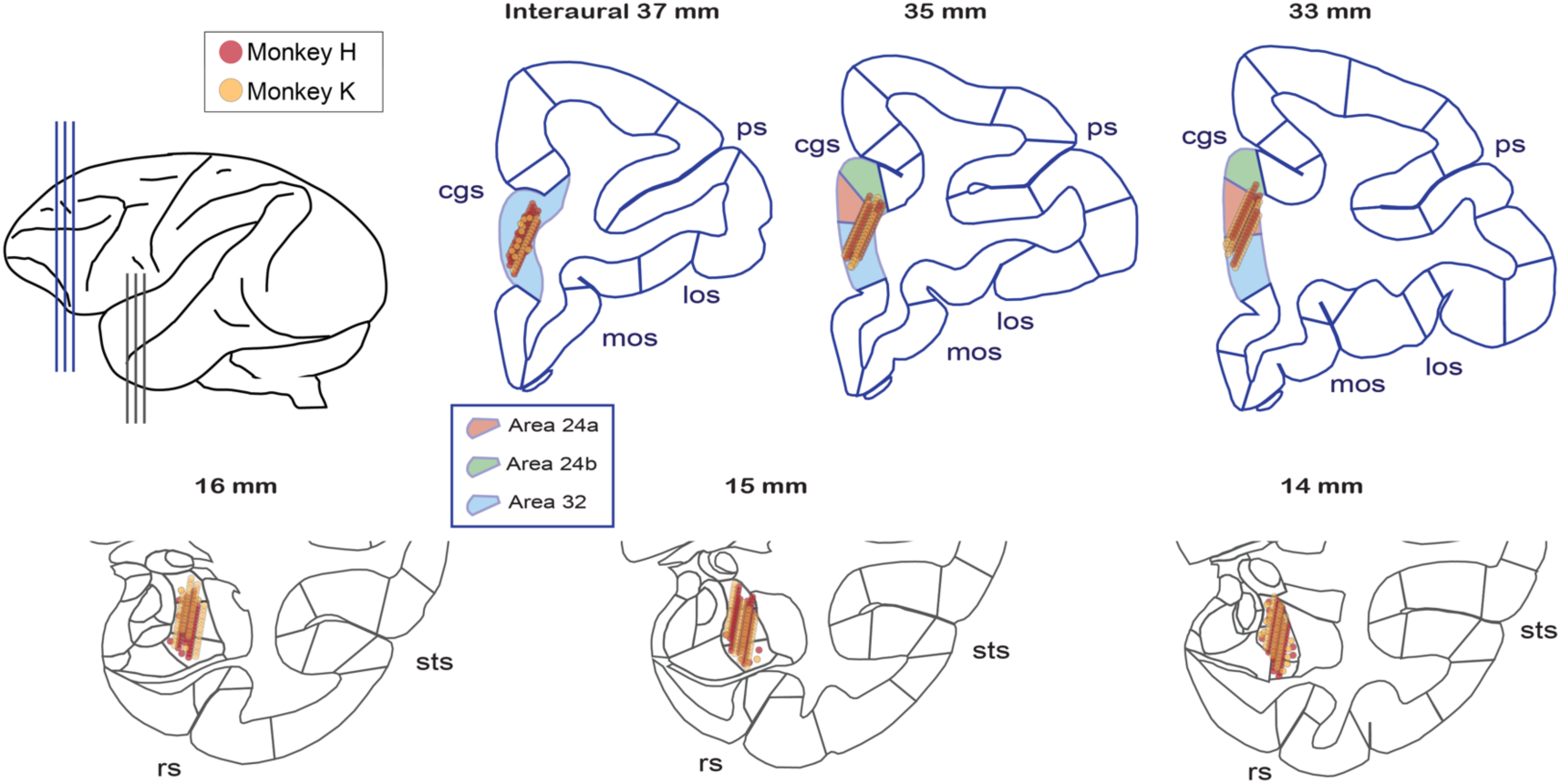
Anatomical locations investigated for the coordination of spiking and LFP activity between BLA and ACCg. Recording locations for individual cells and LFP sites from monkey H (red points) and monkey K (orange points) projected onto the standard stereotaxic coordinates of the rhesus macaque brain atlas^29^. For each area’s projections, three representative coronal slices were chosen with a 2-mm interaural spacing for ACCg and with a 1-mm interaural spacing for BLA in the anterior-to-posterior dimension (as shown in the top left cartoon). Selected landmarks are labeled: cingulate sulcus (cgs), principle sulcus (ps), medial orbitofrontal sulcus (mos), lateral orbitofrontal sulcus (los), superior temporal sulcus (sts), and rhinal sulcus (rs). Boxed inset shows region assignments for the ACC Brodmann names based on the Paxinos atlas^29^.

To determine whether and how neuronal coordination between BLA and ACCg might underlie social decision-making, we related spiking activity of individual cells from each area with LFP oscillations from the other area by calculating spike-field coherence from pairs of neurons and LFP sites^25, 26^. Spike-field coherence values were computed from all recorded cells and LFP sites from which we collected the neural data without any selection criteria. This resulted in 253 ACCg cells paired with 268 BLA LFP sites (ACCg_spike_-BLA_field_) and 90 BLA cells paired with 257 ACCg LFP sites (BLA_spike_-ACCg_field_). In particular, we analyzed coherence patterns during the 150 ms period from the time of acquiring a choice target on free-choice trials (post-decision epoch) and also during the 150 ms period from the central cue onset on forced-choice trials, in order to examine coherence patterns specific to active decisions. Importantly, during this epoch, actors were required to maintain gaze fixation on the target for the duration of the epoch to complete their response, thus removing any eye movement confound and also allowing us to match the timing and gaze-fixation precisely between the free- and forced-choice trials. Most crucially, coherence values were always compared in a relative, reward-matched, fashion (i.e., *Other*–*Bottle* for positive ORP, and *Self*–*Both* for negative ORP) such that any observed coherence differences could not be confounded by actors’ contingency for receiving a juice reward. That is, actors never received rewards in the *Other*/*Bottle* context, but always received rewards in the *Self*/*Both* context, and the use of the *Other*–*Bottle* and *Self*–*Both* contrasts effectively removes any self-reward contingency within the two independent contexts.

Differences in spike-field coherence between expressed positive ORP (choosing *Other* over *Bottle*, *Other*–*Bottle*) and expressed negative ORP (choosing *Self* over *Both*, *Self*–*Both*) exhibited frequency-specific coordination as a function of the area that contributed spikes in the pair. Spikes from BLA cells and the LFP from ACCg (BLA_spike_-ACCg_field_) displayed enhanced coherence in the beta frequency range (defined here as 15–25 Hz) for positive ORP (p < 0.0001, Wilcoxon sign rank) but suppressed coherence in the same band for negative ORP (p < 0.0001) (difference between positive and negative ORPs: p < 0.0001, Wilcoxon sign rank; **Fig. 3a-c** and **Fig. S2**). (**Figure 3a, 3d**, and **3g** show the differences of spike-field coherence values between positive and negative ORPs, whereas **Figures 3b, 3e** and **S2** show spike-field coherence values for each decision preference separately). This enhanced versus suppressed coherence difference was present immediately prior to the time of free-choice decision and lasted until around the time of completing the decision (post-decision epoch). By contrast, in the gamma frequency range (defined here as 45–70 Hz), spikes from ACCg cells and LFP from BLA (ACCg_spike_-BLA_field_) exhibited enhanced coherence, again, for positive ORP (p < 0.0001) but suppressed coherence for negative ORP in the same epoch (p < 0.0001) (difference: p < 0.0001; **Fig. 3d-f**). This coherence difference was also present prior to the time of free-choice decision and lasted until around the time of completing the decision. However, this time course appeared to be lagged in time compared to the BLA_spike_-ACCg_field_ coherence in the beta band (**Fig. 3g**; described in more detail below). Additionally, the differences in spike-field coherence between expressing positive ORP and negative ORP did not change as a function of the temporal progression within a session, for both BLA_spike_-ACCg_field_ coherence (beta band, p > 0.75; gamma band, p > 0.11, linear regression) and ACCg_spike_-BLA_field_ coherence (beta band, p > 0.47; gamma band, p > 0.45).

**Fig. 3.**
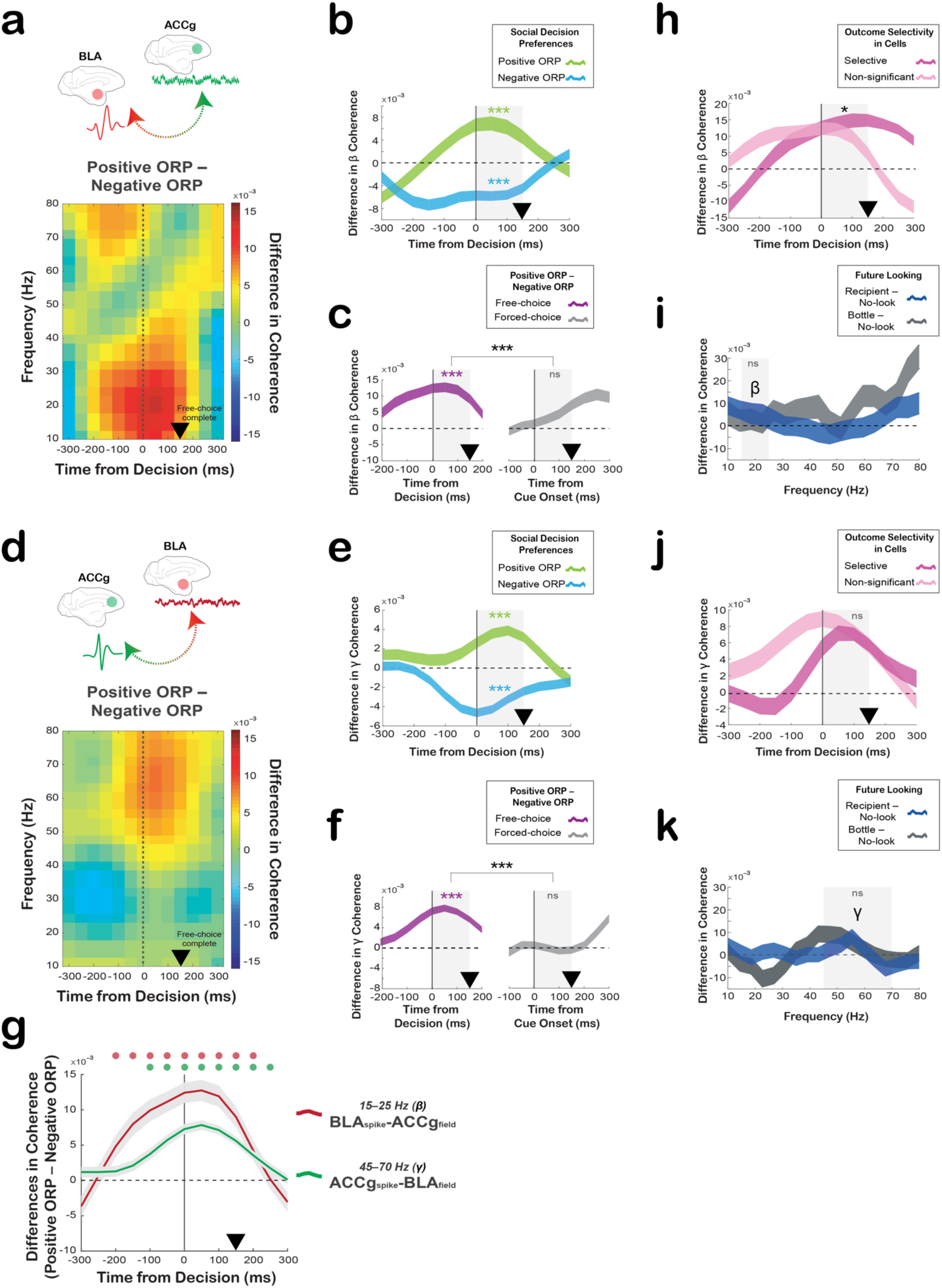
Spike-field coherence between ACCg and BLA shows frequency-specific and free-choice-selective coordination for positive ORP compared to negative ORP. (**a**) Differences in BLA_spike_-ACCg_field_ coherence values between expressing positive ORP (*Other*–*Bottle*) and negative ORP (*Self*–*Both*) across time and frequency aligned to the time of free-choice decision. (**b**) Time courses of the spike-field coherence values in the beta frequency separately for positive ORP (light green; *Other*–*Bottle*) and negative ORP (light blue: *Self*–*Both*). (**c**) Time courses of the beta spike-field coherence differences between expressing positive ORP and negative ORP on free-choice trials (purple) and between the forced-choice construct of positive ORP (*Other-forced*–*Bottle-forced*) and the forced choice construct of negative ORP (*Self-forced*–*Both-forced*) on forced-choice trials (grey). (**d**) Difference in ACCg_spike_-BLA_field_ coherence values between expressing positive ORP and negative ORP across time and frequency. Same format as in a. (**e**) Time courses of the spike-field coherence values in the gamma frequency separately for positive ORP (light green) and negative ORP (light blue). (**f**) Time courses of the gamma spike-field coherence differences between positive and negative ORPs on free-choice (purple) trials and between the forced-choice construct of positive ORP and the forced-choice construct of negative ORP on forced-choice trials (grey). (**g**) Average time courses of the beta band BLA_spike_-ACCg_field_ coherence (red) and the gamma band ACCg_spike_-BLA_field_ coherence (green) differences between the two ORPs. Circles above the lines (in matching colors) show significant differences from zero (p < 0.05, Wilcoxon sign rank). (**h**) Time courses of the spike-field coherence differences between the two ORPs on free-choice trials in the beta frequency separately for outcome selective (dark pink) and non-significant cells (light pink). (**i**) Differences in the BLA_spike_-ACCg_field_ coherence values across frequency between when the monkeys ultimately looked at the conspecific’s face during the inter-trial interval (blue; looking at the conspecific – no-looking) and when they ultimately looked at a bottle (gray; future looking at the bottle – no-looking), collapsed across all outcomes. (**j**) Time courses of the gamma band spike-field coherence differences separately for outcome selective (dark pink) and non-significant cells (light pink) preferences. Same format as **h**. (**k**) Differences in the ACCg_spike_-BLA_field_ coherence values between looking at the conspecific’s face and the bottle in the future. Same format as **i**. In **b-c**, **e-f**, and **h-k**, significant coherence differences from zero (Wilcoxon sign rank) are indicated by asterisks in matching colors and significant coherence differences between traces (Wilcoxon rank sum) are indicated in black asterisks for the analyzed epoch (gray shading) (***, p < 0.0001; **, p < 0.001; *, p < 0.01; ns, not significant). In all plots, the black arrowheads mark the time at which the monkeys completed a free-choice or forced-choice decision by maintaining gaze fixation on a chosen target or cue.

Next, we investigated whether the observed spike-field coherence patterns were stronger for the subsets of BLA and ACCg cells that significantly differentiated decision outcomes (*Self*, *Both*, *Other*, *Bottle*; outcome selective cells) (**Fig. S1**). BLA cells with significant outcome selectivity (37%), exhibited stronger BLA_spike_-ACCg_field_ coherence differences between positive and negative ORPs in the post-decision epoch, compared to the non-significant cells (p < 0.01, Wilcoxon rank sum; **Fig. 3h**). By contrast, ACCg cells with significant outcome selectivity (36%) did not differ in their ACCg_spike_-BLA_field_coherence differences between the two ORPs than the non-significant counterparts (p = 0.12; **Fig. 3j**). These results suggest that outcome-differentiating cells in BLA may play a more specialized role for the BLA_spike_-ACCg_field_ coupling patterns.

Finally, we performed several control analyses to further confirm the enhanced spike-field coupling between BLA and ACCg for expressing positive ORP. We first examined whether the observed spike-field coherence patterns were in any way influenced by actors’ potential intention to look in the future at either the conspecific’s face or the bottle during the inter-trial interval, even though the actors were required to maintain gaze fixation steadily in the main analysis epoch. Specifically, we tested possible differences in spike-field coherence patterns (in all frequency bands) during the post-decision epoch on those trials where the actors ultimately looked at the face (compared to no future looking) as well as those trials where they ultimately looked at the bottle (compared to no future looking). Across all frequency bands examined, we did not observe marked differences. Crucially, we found no differences in the beta band BLA_spike_-ACCg_field_ (p = 0.39, Wilcoxon rank sum; **Fig. 3i**) and the gamma band ACCg_spike_-BLA_field_ coherence (p = 0.77, **Fig. 3k**) patterns, supporting that the observed spike-field coherence cannot be explained by potential anticipatory attentional allocation to the conspecific or the bottle. Second, we ruled out several additional factors from explaining our main findings. The observed spike-field coherence patterns were not simply driven by changes in spiking activity or LFP powers (**Fig. S3** and Supplemental Results, see also **Fig. S4** for LFP power temporal evolution in the beta and gamma bands), or by a more global-level synchrony or common input signals by comparing them to field-field coherence patterns (**Fig. S5** and Supplemental Results). We also examined whether the between-region spike-field coherence patterns reported here were different from the within-region spike-field coherence patterns and found that they were different in several ways (**Fig. S6** and Supplemental Results). Moreover, to test if similar coherence patterns were present even when we construct positive other-regarding and negative other-regarding choices in different ways (type 2 contrasts), we contrasted *Both*–*Self* for delivering rewards to the conspecific and *Bottle*–*Other* for not delivering rewards to the other monkey. We found largely consistent spike-field (**Fig. S7** and Supplemental Results) and field-field coherence patterns with the type 2 contrasts (**Fig. S8** and Supplemental Results), indicating that the spike-field coherence patterns are not the mere product of a preferred choice but are driven by positive other-regarding decisions resulting in other’s rewards. Finally, we ruled out a possibility that sensory-evoked responses associated with choosing a target stimulus might underlie the differential, frequency-specific, coordination between BLA and ACCg. In both beta and gamma frequency bands, the BLA_spike_-ACCg_field_ and ACCg_spike_-BLA_field_ coherence patterns were not at all differentially modulated by the onset of a fixation stimulus (**Fig. S9** and Supplemental Results). Taken together, the current findings support enhanced inter-regional coherence patterns between the two areas associated with expressing positive compared to negative ORP toward a conspecific.

Crucially, the coordination of spikes and LFP observed between BLA and ACCg was specific to when the actor monkeys made preference-based decisions (free-choice). From pseudo-randomly interleaved forced-choice trials in which the computer selected the reward outcomes that were otherwise identical, we were able to construct spike-field coherence differences with matching reward outcomes in the absence of decision-making. We contrasted *Other-forced* and *Bottle-forced* trials (forced-choice construct of positive ORP) for comparing it to positive ORP and contrasted *Self-forced* and *Both-forced* trials (forced-choice construct of negative ORP) for comparing it to negative ORP. The beta band BLA_spike_-ACCg_field_ coherence as well as the gamma band ACCg_spike_-BLA_field_ coherence markedly differed between when the monkeys did or did not make active decisions (**Fig. 3c, f**, and **Fig. S2**). The beta BLA_spike_-ACCg_field_ coherence (15–25 Hz), which was selectively enhanced for positive ORP (p < 0.0001, Wilcoxon sign rank), was absent for the forced-choice positive ORP (p = 0.17) (difference between free-choice and forced-choice: p < 0.0001, Wilcoxon rank sum, **Fig 3c**). Similarly, the gamma ACCg_spike_-BLA_field_ coherence (45–70 Hz), which was again selectively enhanced for positive ORP (p < 0.0001), was absent for forced-choice positive ORP (p = 0.62) (difference between free-choice and forced-choice: p < 0.0001). Therefore, the coordination signatures differentiating positive from negative ORP were unique to making free-choice decisions and not merely driven by either the visual stimuli or the anticipation of specific reward outcomes.

Given that the BLA_spike_-ACCg_field_ coherence differences in the beta band appeared to emerge earlier and terminate sooner than ACCg_spike_-BLA_field_ coherence differences in the gamma band (**Fig. 3**), we next examined possible disparities in the coherence onset time to help elucidate any potential functional differences between the two coordination types. The BLA_spike_-ACCg_field_ coherence in the beta band began to significantly differentiate positive from negative ORP earlier (p < 0.05, Wilcoxon sign rank) than the ACCg_spike_-BLA_field_ coherence in the gamma band (**Fig. 3g**). Additionally, the ACCg_spike_-BLA_field_ coherence in the gamma band continued to significantly differentiate positive from negative ORP longer compared to BLA_spike_-ACCg_field_ coherence in the beta band (**Fig. 3g**). To further investigate the temporal profiles, we examined the time at which either spiking or LFP activity began to significantly signal decision outcomes (**Fig. S10**). Spiking activity associated with choosing *Other* emerged earlier in BLA compared to ACCg (p = 0.001; two sample Kolmogorov-Smirnov test). By contrast, there were no such differences associated with choosing *Self*, *Both*, or *Bottle* outcomes between the two areas (all p > 0.08) (**Fig. S10a**). Further, we did not observe any temporal differences in LFP power between the two nodes for both the beta (*Self*, *Both, Other,* and *Bottle*, all p > 0.38) and the gamma bands (all p > 0.62) (**Fig. S10b**). Finally, we tested if there were any anatomical differences in the strength of spike-field coherence patterns. We found no discernable anatomical gradients for either the beta or gamma spike-field coherence differences between positive and negative ORPs within ACCg and BLA cells/sites (all comparisons using AP, ML, or Depth dimension separately, or based on principal component analysis, all |r| < 0.32, all p > 0.16, Spearman correlation).

### Directionality of information flow between ACCg and BLA for social decisions

Coordination between ACCg and BLA may exhibit a specific directionality of information flow that may critically differ between expressing the two ORPs. To determine this, we performed a partial directed coherence (PDC) analysis, a specialized methodology derived from the Granger analytic principle purposely tailored for analyzing directionality in the frequency-time domain^30^. Without choosing any frequency bands a priori, we observed systematic differences in directional information flow between ACCg and BLA as a function of social decision preference as well as frequency band. We found a significant influence of BLA to ACCg in the beta band (BLA➔ACCg) for positive ORP that began right around the time of decision and continued for the duration of the post-decision epoch (PDC difference between BLA➔ACCg and ACCg➔BLA, p < 0.0001; Wilcoxon sign rank) (**Fig. 4a, b**). This increase in directional influence occurred in the same frequency range that exhibited an increase in the BLA_spike_-ACCg_field_ coherence for positive ORP. By contrast, we found the opposite pattern for negative ORP, with a stronger influence of ACCg to BLA (PDC difference in the beta band between ACCg➔BLA and BLA➔ACCg, p = 0.002). Similarly, we also found a significant but less pronounced influence of BLA to ACCg in the gamma band (BLA➔ACCg) for positive ORP (PDC difference in the gamma band between BLA➔ACCg and ACCg➔BLA, p = 0.04) that appeared later than the BLA➔ACCg influence in the beta band (**Fig. 4c**), with a more robust and again opposite influence of ACCg to BLA for negative ORP (PDC difference between ACCg➔BLA and BLA➔ACCg, < 0.0001). However, while we found frequency-dependent BLA➔ACCg influence for positive ORP in the beta and gamma bands (compared to ACCg➔BLA), the directionality patterns associated with negative ORP were largely frequency-independent between BLA➔ACCg and ACCg➔BLA (**Fig. 4a, b**).

**Fig. 4.**
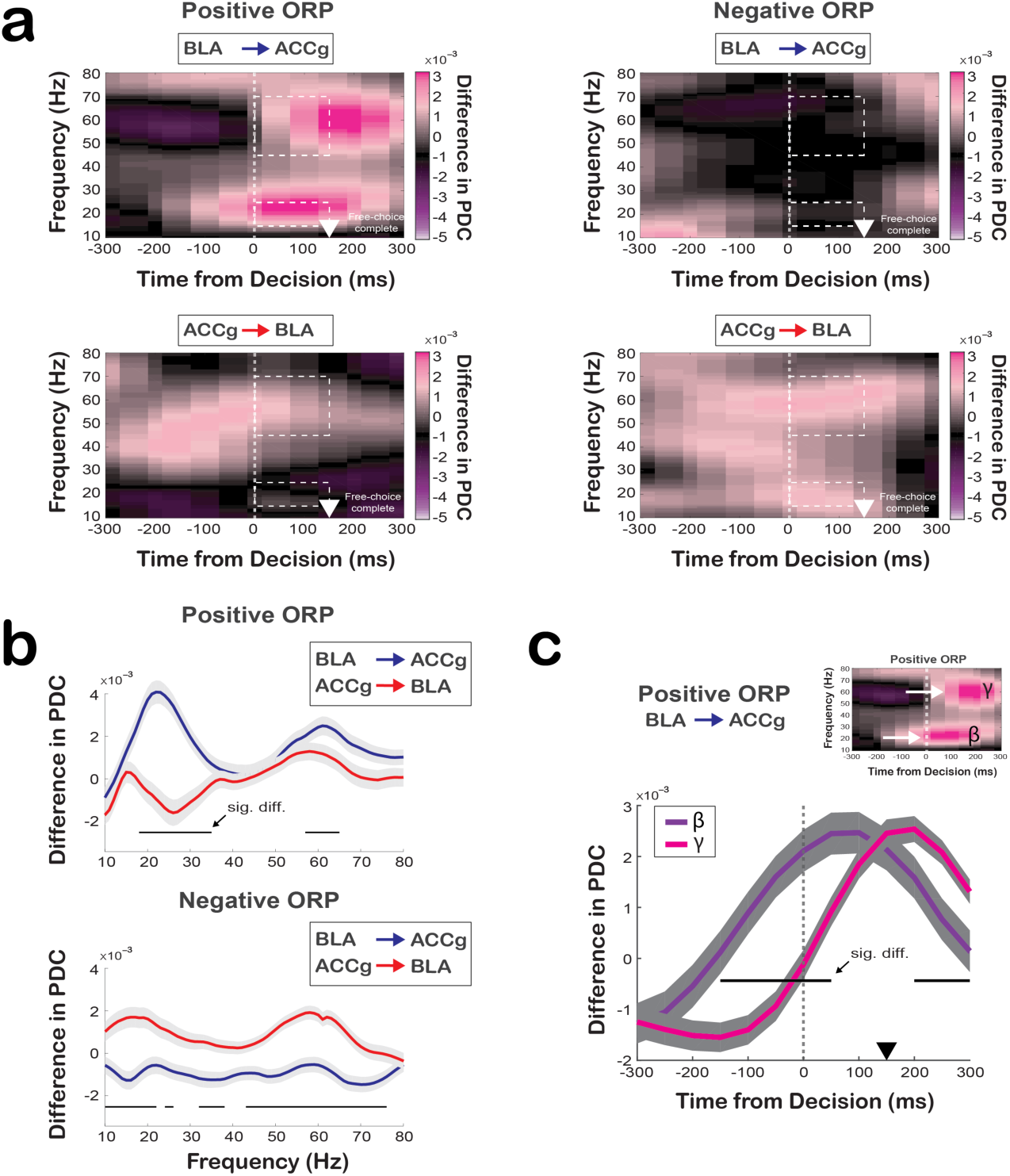
Directionality of information flow between ACCg and BLA for positive ORP and negative ORP as a function of time and frequency. (**a**) Frequency-domain directional influences assessed by partial directed coherence (PDC) on free-choice trials. PDC values as a function of time and frequency for positive ORP (*Other*– *Bottle*) for BLA➔ACCg (top left) and ACCg➔BLA (bottom left), and PDC values for negative ORP (*Self*–*Both*) for BLA➔ACCg (top right) and ACCg➔BLA (bottom right). The white arrowheads mark the time at which the monkeys completed a free-choice by maintaining fixation on a chosen target for 150 ms. Dotted lines indicate the beta (15–25Hz) and gamma (45–70Hz) band during the post-decision epoch. (**b**) Quantification of the directionality of information flow during the free-choice decision epoch as a function of frequency for positive ORP decision (left) and negative ORP (right) for BLA➔ACCg (in blue) and ACCg➔BLA (in red). Horizontal purple lines indicate significantly different between PDC values (p < 0.05, Wilcoxon sign rank). Shaded regions represent standard errors. Horizontal lines indicate significant differences between BLA➔ACCg and ACCg➔BLA (p < 0.05, Wilcoxon sign rank). (**c**) Time courses of the beta and gamma band PDC differences for BLA➔ACCg for positive ORP. Horizontal lines indicate significant differences between the beta and gamma band PDC differences (p < 0.05, Wilcoxon sign rank). Shaded regions represent standard errors.

Finally, we observed similar directionality of information flow in both BLA➔ACCg and ACCg➔BLA for free-choice compared to forced-choice trials for both types of ORPs (**Fig. S11**). While we observed a general BLA➔ACCg influence in the frequency range encompassing both the beta and low gamma bands for positive ORP, the directionality patterns associated with forced-choice trials were much less frequency-dependent compared to the free-choice trials. The directional information flow for negative ORP showed a strong ACCg➔BLA influence (again, opposite to the positive ORP results) for negative ORP with a longer time span.

Together, these findings demonstrate the presence of specific information flow directions between BLA and ACCg, with a general BLA➔ACCg influence for expressing positive ORP and ACCg➔BLA influence for expressing negative ORP, both for free-choice and forced-choice trials. Moreover, even though the PDC analyses do not use spikes, the BLA➔ACCg information flow for positive ORP was observed in the same beta band that exhibited the enhanced BLA_spike_-ACCg_field_ coherence for positive compared to negative ORP.

### Decoding social decisions directly from synchrony between ACCg and BLA

To examine whether neuronal coordination between ACCg and BLA contain decodable information on monkey’s social decisions, we trained a linear decoder to discriminate decision types directly from observed spike-field coherence values (**Fig. 3**). The classifier was trained using randomly selected subsets of 75% of trials and later tested on the remaining 25% of trials used as inputs, yielding estimates of the decision outcome on each trial.

The first decoder was trained to distinguish between *Other* and *Bottle* decisions (positive ORP) from the BLA_spike_-ACCg_field_ coherence values in the beta band (15–25 Hz) or from the ACCg_spike_-BLA_field_ coherence values in the gamma band (45–70 Hz) across time. Decoding performance from the beta BLA_spike_-ACCg_field_ coherence for discriminating *Other* from *Bottle* began to increase prior to the decision time and peaked around the time of the decision (p < 0.0001, compared to an empirically-derived null distribution, Wilcoxon sign rank) (**Fig. 5a**). On the other hand, the decoding accuracy from the gamma ACCg_spike_-BLA_field_ coherence for discriminating *Other* from *Bottle* was lower at the time of free-choice decision but gradually improved during the post-decision epoch as monkeys fixated on a chosen option to complete the decision (**Fig. 5b**). The second decoder was trained to distinguish between *Self* and *Both* for classifying negative ORP in the identical frequency bands and times. Compared to the first decoder, the decoding performance was overall lower (positive vs. negative ORP in the post-decision epoch: p < 0.0001 and p < 0.0001 for decoding from the BLA_spike_-ACCg_field_ and ACCg_spike_-BLA_field_ coherence, respectively) and did not show clear time-locked increases around the time of free-choice decision, albeit still being able to decode above its empirically-derived chance level (**Fig. 5a**, **b**). In order to establish whether improved decoding performance for positive ORP might emerge earlier in BLA_spike_-ACCg_field_ compared to ACCg_spike_-BLA_field_ coherence patterns, we divided the analysis window into the first and second halves of the post-decision epoch and directly compared decoder performance between the two ORPs in each period (beta BLA_spike_-ACCg_field_ vs. gamma ACCg_spike_-BLA_field_). Consistent with this hypothesis, decoding performance was significantly greater for the BLA_spike_-ACCg_field_ in the beta band compared to the ACCg_spike_-BLA_field_ coherence in the gamma band in the earlier phase (p < 0.0001), whereas this relationship was reversed in the later phase of the epoch, such that relative decoding performance for the ACCg_spike_-BLA_field_ in the gamma band was significantly greater than the BLA_spike_-ACCg_field_ coherence in the beta band (p < 0.0001) (**Fig. 5c**). These temporal differences in decoding accuracy were consistent with the temporal differences observed between the beta BLA_spike_-ACCg_field_ and the gamma ACCg_spike_-BLA_field_ coherence differences in favor of positive ORP. Overall, although the extent of decoding accuracy for predicting monkey’s social decisions was low even at the peak accuracy level, decoding directly from the synchrony signatures was nevertheless reliable.

**Fig. 5.**
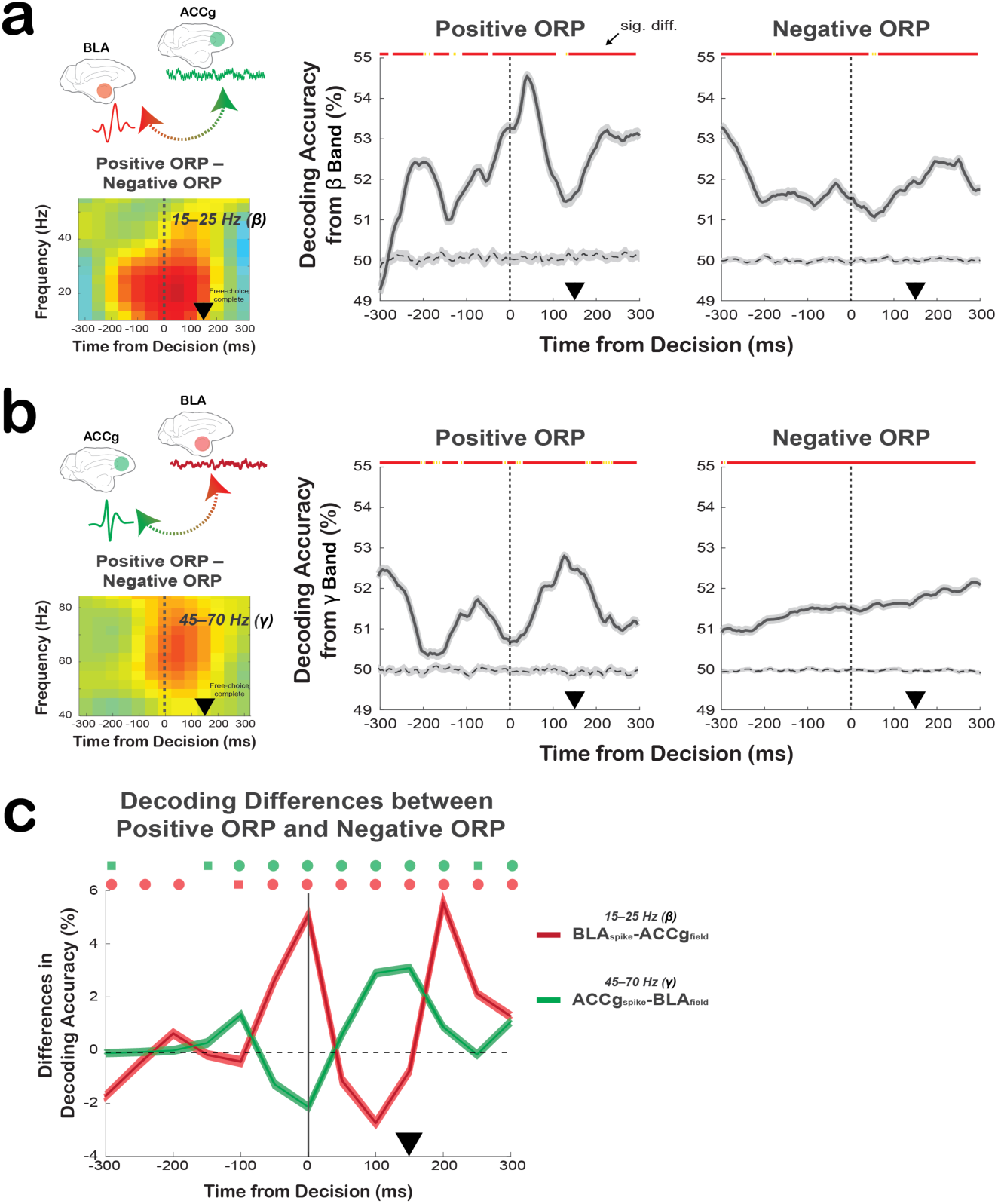
Decoding social decisions directly from the spike-field relations between ACCg and BLA. (**a**) Decoding performance using the BLA_spike_-ACCg_field_ coherence differences in the beta band (shown in the left inset) for discriminating *Other* from *Bottle* (middle) and discriminating *Self* from *Both* (right) decisions over time (mean ± s.e.m.). Dashed lines represent empirically determined null distribution. (**b**) Average decoding performance using the ACCg_spike_-BLA_field_ coherence in the gamma band (shown in the left inset) for discriminating *Other* from *Bottle* (middle) and discriminating *Self* from *Both* (right) decisions over time. Same format as in **a**. In **a** and **b** lines indicate significant differences from the null in each of the 5 ms bin (red: p < 0.0001, yellow: p < 0.05, Wilcoxon sign rank). (**c**) Differences in decoding performances between *Other*/*Bottle* and *Self*/*Both* contexts from the beta BLA_spike_-ACCg_field_ coherence (red) and the gamma ACCg_spike_-BLA_field_ coherence (green). Symbols above the lines (in matching colors) show significant differences from zero (circles: p < 0.0001, square, p < 0.05; Wilcoxon sign rank). In all plots, the black arrowheads mark the time at which the monkeys completed a free-choice decision by maintaining fixation on a chosen target for 150 ms.

## Discussion

Coordination through oscillatory mechanisms has been theorized to provide a unique temporal window for neuron-to-neuron synchrony^31–33^. Growing evidence supports that oscillatory coordination across different brain regions is one mechanism used to regulate a wide range of cognitive functions, from visual perception^34, 35^, motor planning^36^, and spatial navigation^37^ to higher-order functions underlying working memory^38^, associative learning and decision-making^39–42^. A number of studies have also emphasized the importance of cortical-subcortical interactions in facilitating complex cognitive operations^20, 21, 24, 40–42^. The frequency-specific and direction-selective coordination between BLA and ACCg reported here exemplifies one possible medial prefrontal-amygdala coordination mechanism by which two nodes in the social brain network interact during complex social behaviors.

The coherence patterns between ACCg and BLA were predominantly characterized by enhanced coherence for positive ORP but suppressed coherence for negative ORP (**Fig. 3**). Thus, enhanced co-engagements of ACCg and BLA may promote the expression of positive ORP, whereas co-disengagements of ACCg and BLA in turn may lead to the expression of negative ORP. Notably, the coordination patterns exhibited specializations in the frequency domain. Our results suggest that the beta band may be involved in linking spiking outputs from BLA cells with the synaptic input or dendritic integration of ACCg cells, whereas the gamma band may be involved in the interaction linking these processes in reverse. Frequency-specific coordination between ACCg and BLA may provide separate synchrony “streams” that could be useful in mediating processes related to social decision preference. Such specializations of frequency channels underlying different cognitive operations have also been observed in the past for cortico-cortical interactions involving top-down and bottom-up visual attention^43^.

In general, synchrony found in lower frequency range is thought to be more robust to temporal dynamics of spiking activity due to slower temporal profiles^44^, perhaps making lower frequency channels better for synchronizing distant structures. Further, the beta frequency in particular has been theorized to mediate ‘the status quo’ functions associated with maintaining predicted or internally-consistent behaviors^45^. The use of a beta frequency channel for linking BLA spikes to ACCg field may be one mechanism for facilitating robust and long-range coordination between BLA and ACCg for positive ORP. On the other hand, synchrony found in higher frequency range is likely be strongly driven by local computations, requiring fast-spiking GABA-ergic inhibitory interneurons^44, 46^. The gamma frequency range especially has been associated with generating selective representations of certain stimuli over others^47^. The use of a gamma frequency channel for linking ACCg spikes to BLA field may be reflective of further interactions based on local computations in ACCg following the long-range synchrony initiated through the beta frequency.

Importantly, the directionality of information flow between the two regions was largely selective for positive ORP, with the predominant directional influence from BLA to ACCg in the beta frequency channel greater for positive compared to negative ORP. This directionality occurred in the same frequency band that exhibited enhanced coordination between BLA spikes and ACCg field for positive ORP. Moreover, the BLA_spike_-ACCg_field_ coordination associated with positive ORP was amplified for the outcome selective BLA cells. Taking these results together with earlier emergence of the BLA_spike_-ACCg_field_ compared to the ACCg_spike_-BLA_field_ coordination, spiking activity from BLA cells that differentiate social decision outcomes may drive ACCg for processing positive ORP. BLA cells are well-known for signaling social contextual information, such as distinct social gaze orientations and facial expressions^48, 49^, that powerfully shapes social behaviors. Future work can test if and when BLA cells with other specialized functions transmit such information to rostral ACCg or other medial prefrontal cortical areas to bias social decisions across various social contexts.

Interestingly, the coordination between ACCg and BLA was largely specific to free-choice or active decisions, compared to trials on which the computer made the decisions for the actors. This finding argues that the coordination between these areas was not driven by anticipation of upcoming reward outcomes, but rather by voluntarily expressing one’s social decision preferences. Although it is inherently difficult to entirely rule out the possibility that these circuits are simply less engaged by virtue of not making active decisions on forced-choice trials, freely expressing social preference may engage the medial prefrontal-amygdala circuit in unique ways. This hypothesis is also supported by two previous observations in the primate BLA demonstrating specialized neural codes for computing free-choice, compared to forced-choice, decisions^9, 50^.

In social decision-making scenarios like the one abstracted by our task, it is imperative for a decision-maker to be aware of a chosen option and an ultimate actualization of the corresponding reward outcome for either self or other. In the reinforcement learning theory, post-decision or ‘afterstate’ signals available during post-decisional monitoring can serve as an important and unique feedback mechanism for more efficient learning of actions and reward outcomes^59^. We hypothesize that the specialized coordination of BLA and ACCg prioritizing positive ORP during the post-decision state may indicate that the two regions coordinate to synchronize post-decision processing for efficiently linking across action, chosen value, and the ultimate reward outcome of another individual. However, future work with a specific behavioral design for modulating the fidelity of post-decision monitoring in relation to BLA-ACCg coupling is necessary to more directly test this hypothesis.

Finally, it is worth pointing out some limitations of the current work. Although the task had an embedded condition for delivering juice to a non-social entity (bottle), it remains unknown whether similar coherence patterns would be present when expressing a preference in a completely non-social context. Future work should examine how the reported spike-field coherence patterns between BLA and ACCg might be differentially modulated by expressing decision preferences in social and non-social contexts. Moreover, despite the fact that we removed any self-reward contingency within the two independent decision-making contexts (*Self*–*Both* from *Self*/*Both* context and *Other*–*Bottle* from *Other*/*Bottle* context), it is worthwhile to acknowledge that the two contexts were clearly different and deriving positive ORP from *Other*/*Bottle* context and negative ORP from *Self*/*Both* context might have influenced our findings. However, the fact that we observed overwhelmingly similar spike-field as well as field-field coherence patterns upon deriving positive ORP from the *Self*/*Both* context (*Both*–*Self*) and negative ORP from the *Other*/*Bottle* context (*Bottle*–*Other*) greatly mitigates this concern.

Overall, the current findings support the view that BLA and ACCg neurons utilize distinct frequency channels and direction-selective coordination in social decision-making. Efficient and perhaps even strategic coordination occurring between medial prefrontal regions and the amygdala that prioritizes positive ORP over negative ORP may play an essential role in promoting mutually beneficial social cohesion. In turn, failures in synchronized transmissions along the medial prefrontal-amygdala network may bias other relevant brain networks to converge toward producing atypical social behaviors.

## Data Availability

Behavioral and neural data presented in this paper and the main analysis codes will be available through https://github.com/changlabneuro upon acceptance of the manuscript.

## Acknowledgements

We are extremely grateful to Bijan Pesaran for his guidance on examining oscillatory neural processes throughout the duration of this research. We especially thank Daeyeol Lee and Alex Kwan for their thoughtful discussions and suggestions on improving this work. We also thank Amrita Nair and Siqi Fan for insightful comments on the manuscript. This work was supported by the National Institute of Mental Health (R01MH110750; R01MH120081; R21MH107853; R00MH099093), Alfred P. Sloan Foundation (FG-2015-66028), and the Teresa Seessel Postdoctoral Fellowship.

## Author Contributions

S.W.C.C. and O.D.M. designed the study and wrote the paper. O.D.M. performed the experiments.

C.J.C., N.A.F., O.D.M., and S.W.C.C. analyzed the data.

## Competing Financial Interests

The authors declare no competing financial interests.

## Online Methods

### Animals

Two adult male rhesus macaques (*Macaca mulatta*) were involved in the study as actors (monkeys K and H; ages, both 6; weights, 7 and 8 kg), and two adult female monkeys (ages, 6 and 10; weights, 9 and 10 kg) were involved only as recipients in the social reward allocation task. All animals were unrelated and not cagemates. Actors were housed in a colony room with other male macaques, whereas two female macaques resided in an adjacent colony room with other females. All four subjects were housed in pairs with other animals from the colony, kept on a 12-hr light/dark cycle, had unrestricted access to food, and controlled access to fluid during testing. All procedures were approved by the Yale Institutional Animal Care and Use Committee and in compliance with the National Institutes of Health Guide for the Care and Use of Laboratory Animals.

### Surgery and anatomical localization

All four animals received a surgically implanted headpost (Grey Matter Research) for restraining their head during the experiments. Subsequently, a second surgery was performed on actor monkeys to implant a recording chamber (Crist) to provide access to ACCg and BLA. Placement of the chambers were guided by both structural magnetic resonance imaging (MRI, 3T Siemens) scan and stereotaxical coordinates. Prior to starting the recording experiments, we performed a manganese (Mn)-enhanced magnetic resonance imaging (MEMRI) session for each actor monkey to precisely localize our recording sites in both ACCg and BLA. For MEMRI, we focally infused 2 μl of 19.8 μg/μl of Mn (manganese (II) chloride) in saline solution in both areas using modified Hamilton syringes that traveled along the identical trajectory as our electrodes. We then performed a structural MRI scan 3 hours after the infusion to visualize a bright halo to confirm anatomical locations^51^. All electrophysiological recordings were carried out simultaneously from ACCg (Brodmann areas 24a, 24b, and 32)^29^ and BLA^29^ (**Fig. 2**).

### Social reward allocation task

Two monkeys (an actor and a recipient) sat in primate chairs (Precision Engineering, Inc.) at 100 cm from one another at a 90° angle (**Fig. 1a**). Each monkey had his own monitor, which displayed identical visual stimuli. Both monkeys had their own juice tubes from which juice drops were delivered via solenoid valves. A third juice tube with its own dedicated solenoid valve delivered juice rewards into an empty bottle (*Bottle*), which was placed on the opposite side of the recipient (**Fig. 1a**). To prevent monkeys from forming secondary associations of solenoid clicks, the three solenoid valves were placed in another room and white noise was played in the background during all experimental sessions. An infrared eye-tracking camera (EyeLink 1000, SR Research) continuously recorded the horizontal and vertical eye positions from actor monkeys.

An actor began a trial by fixating on a central square for 150 ms with gaze. The reward value at stake on each trial was specified by a magnitude cue displayed as a vertical bar indicating juice volume (0.2, 0.4, or 0.6 ml). The actor was required to maintain gaze fixation on the magnitude cue for 400 ms. Following a variable delay (200, 400, or 600 ms), the actor was presented with either a free-choice (75%) or a forced-choice (25%) trial. On free-choice trials, two visual targets appeared at two random peripheral locations on opposite sides of the screen. The actor had 2 sec to make a choice by shifting gaze to a target and maintaining the fixation on the target for additional 150 ms in order to complete a choice (i.e., any break in gaze fixation resulted in an incomplete trial with no further progression into the trial). These choice targets were always presented in two distinct contexts presented pseudo-randomly. In the *Self*/*Both* context (50% of free-choice trials), the actor made decisions to deliver a juice drop to himself (*Self*) or both himself and the recipient monkey (*Both*; the same amount was delivered at the same time to both monkeys). By contrast, in the *Other*/*Bottle* context (50% of free-choice trials), the actor made decisions to deliver a juice drop to the recipient monkey (*Other*) or to the empty juice collection bottle (*Bottle*). Critically, any choice made in the two contexts were ‘reward-matched’ from actor’s perspective such that the actor always received a reward in the *Self*/*Both* context but never received a reward in the *Other*/*Bottle* context. After a following variable delay from completing the decision (200, 400, 600, or 800 ms), a juice reward corresponding to the chosen target was delivered to himself (*Self*), to the recipient (*Other*), to both monkeys (*Both*), or to the bottle (*Bottle*). On forced-choice trials, only a single central cue was presented on the screen, and the actor had to simply maintain the fixation for 150 ms to complete the forced-choice decision (i.e., any break in fixation resulted in an incomplete trial with no further progression into the trial). These computer-determined reward outcomes occurred with equal frequency, pseudorandomly ordered. After a following variable delay (200, 400, 600, or 800 ms), a juice reward corresponding to the central cue was delivered to himself (*Self-forced*), to the recipient (*Other-forced*), to both monkeys (*Both-forced*), or to the bottle (*Bottle-forced*). For both free-choice and forced-choice trials, reward delivery was followed by a 2.5 sec inter-trial interval, during which the actor was free to look at the recipient or any other locations in the setup. A trial was considered incomplete if the actor failed to choose a target or maintain the required 150 ms fixation on free-choice trials or to maintain the required 150 ms fixation on the cue on forced-choice trials. The incomplete trials were not included in the analyses.

### Electrophysiology

LFP and spiking activity was recorded using 16-channel axial array electrodes (U- or V-Probes, Plexon) or single tungsten electrodes (FHC Instruments) placed in each of the recording regions using a 32-channel system (Plexon). At the beginning of each session, a guide tube was used to penetrate the intact dura and to guide electrodes, which were lowered using a motorized multi-electrode microdrive system (NaN Instruments) with a speed of 0.02 mm/sec. After the electrodes reached the target sites in both ACCg and BLA, we waited 30 min for the tissue to settle before starting each recording session to ensure signal stability. Because some of the data were obtained using two 16-channel electrode arrays, one in ACCg and the other in BLA (20% of the total recording sessions), we randomly assigned 16 uniquely paired LFP sites across the two regions, using a random number generator, to remove redundant inflations of correlation for the relevant data.

## Data Analysis

### Behavioral analyses

We constructed a choice preference index as contrast ratios^6, 27, 28, 52^ (Eq. 1).

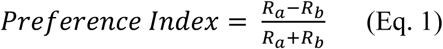

*R_a_* and *R_b_* were the frequency of particular choices. For the *Self*/*Both* context, *R_a_* and *R_b_* were numbers of *Both* and *Self* choices, respectively. For the *Other*/*Bottle* context, *R_a_* and *R_b_* were numbers of *Other* and *Bottle* choices, respectively. An index of 1 thus corresponds to always choosing a positive ORP outcome, –1 corresponds to always choosing a negative ORP outcome, and 0 indicates indifference. We additionally performed a regression analysis to quantify changes over time in their behavioral preferences for both *Self*/*Both* and *Other*/*Bottle* context in each session.

Looking frequency was computed based on the average number of gaze shifts landing on the face of the recipient monkey (the face region of the recipient was empirically mapped and fitted with a rectangle window) or the bottle (mapped empirically with the same-dimensioned window as the face region) during the 2.5 sec inter-trial interval^6, 27, 28, 52^. Decision reaction time, the time from the onset of two targets on free-choice trials to eye movement onset, were computed using a 20° sec^−1^ velocity criterion^6, 27, 28, 52^.

### Spiking and LFP activity

Broadband analog signals were amplified, band-pass filtered (250 Hz–8 kHz), and digitized (40 kHz) using a Plexon OmniPlex system. Spiking data were saved for waveform verifications offline and automatically sorted using the MountainSort algorithm^53^. LFP data were analyzed using custom MATLAB scripts (The MathWorks) and the Chronux signal processing toolbox^54^. Continuous LFP signals from each recording electrode in each area were segmented into 1-sec periods centered on acquiring (i.e. saccade offset) the choice target or acquiring the central cue at a sample rate of 1 kHz. Raw signals were then band-passed filtered from 2.5 Hz to 250 Hz. We chose a zero-phase filter to avoid introducing phase-distortions to the signals. Signals were normalized by subtracting a reference voltage trace recorded from an independent reference electrode placed in the subdural space in order to eliminate the common noise from each electrode. Three primary epochs were used to carry out neural data analyses: during the 150 ms window during the first fixation period required to begin each trial (baseline epoch); during the 150 ms period from the time of acquiring (i.e. saccade offset) a choice target on free-choice trials (post-decision epoch) and also during the 150 ms period after the central cue onset on forced-choice trials (cue epoch). To determine outcome selective cells from each region, we performed one-way ANOVA with outcome as the factor (*Self*, *Both*, *Other*, *Bottle*) using the spiking activity from either the post-decision epoch or reward epoch (50–450 ms from reward onset). Finally, to compare the emergence times of outcome selective signals in both spiking and LFP activity, we calculated the cumulative distributions of the times at which each cell or LFP site exhibited significant encoding of different outcomes around the time of decision-making, relative to the baseline epoch (p < 0.05, Wilcoxon sign rank).

### Spike-field coherence and field-field coherence

We quantified spike-field coherence level by examining the phase differences between LFP and spike signals. We designated one area as the “spike contributor” and the other area as the “field contributor”. Spike-field coherence was calculated from two directions, either ACCg or BLA as the spike contributor and the other area in the pair as the field contributor. We first binned spikes and LFP using sliding time windows of 150 ms, in steps of 50 ms, for a 1 sec interval centered on the time of decision on free-choice trials or the cue onset on forced-choice trials. Fourier estimates were then computed by means of a multi-taper transformation applied to single trial data; we selected a time half-bandwidth product of 2, and multiplied the raw signals by 3 Slepian (orthogonal) tapers^55^. With a 1 kHz sampling rate, this yielded a frequency resolution of ∼3.096 Hz. Spectral density estimates were additionally restricted to the 10–80 Hz interval, considering the Nyquist limit. The spectrum density of point process (spikes) was transformed by applying fast Fourier transform on the discrete data. Coherence was then calculated between two spectrum densities of continuous process (LFP) and point process (spikes) by computing the cross-spectral density of the two processes (*x* and *y; P_xy_*) with respect to frequency (*f*), which was normalized by the product of the power spectral densities of each process (*P_xx_* and *P_yy_*) as a function of frequency (Eq. 2).

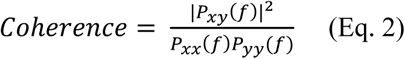

Raw coherence values therefore ranged from 0 to 1, where a perfectly constant phase relationship between the two regions would be indicated by a coherence value of 1 while an absence of any phase relationship would be indicted by a value of 0. We contrasted coherence values between different conditions and obtained average across pairs of cells and LFP sites. A linear regression was used to quantify the changes in BLA_spike_-ACCg_field_ coherence and ACCg_spike_-BLA_field_ coherence patterns for both the beta and gamma band over time within each session.

For calculating within-region spike-field coherence, we used the same approach described above for between-region spike-field coherence but excluded relating spikes and LFPs originating from the same electrode channels. For looking at the relationships of LFPs across the two regions, field-field coherence was computed in the same format as in the spike-field coherence described above except the following. Field-field coherence was calculated between two spectrum densities of continuous processes (LFPs from each region) by computing the cross-spectral density of the two processes (*x* and *y; P_xy_*) with respect to frequency (*f*), which was normalized by the product of the power spectral densities of LFP processes from each region (*P_xx_* and *P_yy_*) with respect to frequency (same format as in Eq. 2).

### Directionality of information flow

We calculated partial directed coherence (PDC), which is based on multivariate autoregressive (MVAR) model and is well suited for describing directionality of information flow between simultaneously recorded time series in the frequency domain^30^. We contrasted time-varying PDC as (*Other*) – (*Bottle*) and (*Self*) – (*Both*) for free-choice trials, as well as (*Other-forced*) – (*Bottle-forced*) and (*Self-forced*) – (*Both-forced*) for forced-choice trials. As we did for the coherence analyses, we restricted the combinations of pairs to be unique across sites. For example, for the data recorded from a 16-channel array placed in each of the two areas, we randomly selected 16 unique pairs out of 16 x 16 pairs to avoid redundancy and undesired inflation in correlations. For each pairwise LFP signals, the parameters of multivariate autoregressive model (MVAR) of order *r* was formulated as:

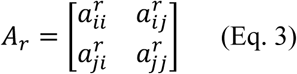

where paramete *a* reflects linear relationship between channel *i* and *j* at delay *r*. While *r* = 1 … *p* represents the order of the model. To obtain PDC measures across time, instead of applying adaptive filtering method to estimate time-varying autoregressive coefficient, we calculated PDC values based on sliding window of 150 ms with a 50 ms step size just as we do for the coherence measures. Model order of MVAR model was estimated by using the post-decision epoch data to minimize Schwarz Bayesian information criteria (SBC) across all LFP pairs. This resulted in p = 12, specifying that the current value is predicted by immediately preceding twelve values in the series. The model extended to the frequency dimension was defined as:

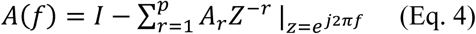

where *I* is the identity matrix and *f* ranges within 0 to Nyquist frequency. PDC values were then defined by taking the absolute value of *A(f)* and normalizing by its column vector (see equation 18 in reference 30). To reduce the co-variability of signal between channels due to common sources, we adapted the extended version of classical PDC^57^. The new generalized orthogonalized measure of PDC 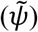 as a function of time and frequency was defined as:

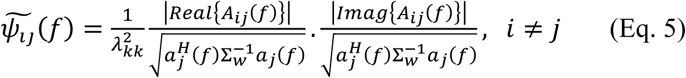

where *a_j_* is the *j*’s column vector and *A_ij_* is the *ij*th element of *A*(*f*). *H* denotes the Hamilton transpose of the vector *a*. Σ*_w_* is the diagonal covariance matrix from MVAR noise covariance matrix *w*, where λ*_kk_* is a diagonal element of Σ*_w_*. For one pair of channels, 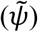 was shown in a 2 x 2 matrix, where non-diagonal elements represent directional interaction between channel *i* and *j*, that is, ACCg➔BLA or BLA➔ACCg. We then calculated and averaged 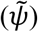 across all trials in each condition (*Self*, *Both*, *Other*, or *Bottle*) and averaged pairwise sites of PDC across all recording sessions. For testing whether specific frequency bands exhibit significantly different PDC values between conditions for each ACCg➔BLA and BLA➔ACCg, we compared PDC values from the same time window used for the main spike-field coherence results.

### Linear Discriminant analysis (LDA)

To test the decodability of social decisions directly from spike-field coherence values, we used a standard linear classifier for population decoding^58^. The analysis was run separately for each time-frequency bin (150 ms bin with 5 ms steps) and for each decision context. For a given time-frequency bin and context, the trial-level vector of spike-field coherence values in that bin was extracted, along with the corresponding vector of decision outcomes for each trial. This outcome vector contained *Other* and *Bottle* labels or *Self* and *Both* labels, depending on the decision context. The decoder was therefore trained to discriminate between binary outcomes on the basis of spike-field coherence values. In the training phase, 75% of trials were selected at random to train the classifier model. In the testing phase, coherence values for the remaining 25% of trials were used as inputs, yielding estimates of the decision outcome on each trial.

Decoder performance was assessed as the percentage of test-phase trials that were correctly labeled. The statistical significance of the performance was assessed with a permutation test. For each of 100 iterations, a null value of the decoder’s performance was obtained by shuffling the decision outcome labels before training and testing. The analysis thus produced arrays of matching sizes representing the real and null decoding performance for each (time, frequency, condition, iteration) sequence. Decoding was considered significant if the average performance was higher than the corresponding null performance at least 99% of the time (p < 0.01, FDR-corrected for multiple comparisons across frequencies).

## Supplementary Results

### Absence of correlations between spike-field coherence and the magnitudes of firing rates or LFP power

To test whether the differences observed in spike-field coherence could have been driven simply by either the changes in spiking activity or LFP power *within* each brain region, we correlated the changes in firing rate or the changes in LFP power of each brain region with the observed BLA_spike_-ACCg_field_ and ACCg_spike_-BLA_field_ coherence patterns (**Fig. S3**). High, medium, and low magnitude quantiles from the firing rates of BLA cells were not correlated with the beta band BLA_spike_-ACCg_field_ coherence values between the two ORPs (post-decision epoch, r = –0.12, p = 0.43, Pearson correlation). Similarly, the firing rates of ACCg cells also did not relate to the gamma band ACCg_spike_-BLA_field_ coherence values (r = 0.05, p = 0.53) (**Fig S3a**). Moreover, we found a similar pattern of results for LFP powers; the changes in the beta band BLA_spike_-ACCg_field_ coherence had only trending relations with the beta band LFP power in ACCg (r = 0.10, p = 0.05) and changes in the gamma band ACCg_spike_-BLA_field_ coherence were not correlated with the gamma band LFP power in BLA (r = –0.03, p = 0.23) (**Fig. S3b**) (also see **Fig. S4** for the time courses of LFP power in beta and gamma bands).

### Field-field coherence between BLA and ACCg does not account for spike-field coherence patterns

To establish if the observed spike-field coherence patterns were not merely driven by a more global-level synchrony or by common input signals, we examined field-field coherence values across the two areas, which are known to be more susceptible to such non-specific sources, in order to compare them to the spike-field coherence values (**Fig. S5c-d**).

Field-field coherence patterns were examined from 447 ACCg sites (241 and 206 sites in monkey H and K, respectively) and 402 BLA sites (211 and 191 sites in monkey H and K). During the post-decision epoch, we found no increase in the field-field coherence patterns in the beta band for positive ORP (p > 0.34, Wilcoxon sign rank) and also no decrease in the field-field coherence patterns for negative ORP (all p > 0.21) (**Fig. S5a**). Rather, we observed significantly enhanced field-field coherence values in the gamma band for negative ORP (p < 0.0001), without any changes in the gamma band associated with positive ORP (p > 0.19). We next compared the field-field coherence values between free-choice and forced-choice trials. We found a significant difference in the beta band field-field coherence between free- and forced-choice trials (p < 0.0001, Wilcoxon rank sum, **Fig S5b**), with a slight increase associated with free-choice and a strong decrease for forced-choice. In the gamma band, however, we did not observe any differences between the two trial types (p > 0.51).

When we directly compared the spike-field coherence differences to the field-field coherence differences between positive and negative ORPs, we found a markedly stronger spike-field coherence patterns both in the beta and in gamma bands (post-decision epoch, both p < 0.0001, Wilcoxon rank sum) (**Fig. S5c**). Therefore, the observed spike-field coherence patterns differentiating positive ORP from negative ORP (**Fig. 3**) were not likely to be simply driven by common input signals to both ACCg and BLA, as these should be better indexed by their field-field patterns.

### Within-region spike-field coherence in BLA or ACCg does not account for between-region spike-field coherence patterns

To identify potential relationships between within-region and between-region coherence patterns, we compared the interareal spike-field coherence patterns directly to the within-region spike-field coherence patterns (**Fig. S6**). Differences in BLA_spike_-BLA_field_ coherence between positive and negative ORPs increased diffusely before the time of decision not only in the beta band but also in a much wider gamma range (both frequency bands, both p < 0.0001, Wilcoxon sign rank; **Fig. S6a**). There were overall similar coherence differences on forced-choice trials when the computer determined the outcomes (both bands, p < 0.0001) (**Fig. S6b**). On the other hand, differences in ACCg_spike_-ACCg_field_ coherence between positive and negative ORPs increased around the time of decision in the gamma band (both p < 0.0001; **Fig. S6a**). This coherence difference was not present in the absence of decision-making on forced-choice trials (p = 0.28; **Fig. S6b**).

Next, we directly compared the within-region and the between-region spike-field coherence patterns. The ACCg_spike_-ACCg_field_ coherence differences between the two ORPs were substantially weaker compared to the BLA_spike_-ACCg_field_ coherence differences in the beta band (p < 0.0001, Wilcoxon rank sum; **Fig. S6c-d**). Moreover, although the BLA_spike_-BLA_field_ coherence differences between the two ORPs were more comparable to the ACCg_spike_-BLA_field_ coherence differences in the gamma band (p = 0.12), the peak of the within-region coherence differences occurred prior to the time of choice and these values were in the process of decreasing toward the baseline during the post-decision epoch (**Fig. S6c-d**). Therefore, although the within-region spike-field coherence patterns showed interesting and systematic effects, there were several notable differences compared to the between-region spike-field coherence patterns (**Fig. 3**).

### Different ways of contrasting positive and negative ORP replicate the main coherence findings

To investigate whether the original spike-field and filed-field coherence findings were not due to the directions of the contrasts we chose, we performed the identical coherence analyses (spike-field coherence, **Fig. S7** and field-field coherence, **Fig. S8**) for the following additional contrasts: *Both*–*Self* and for *Bottle*–*Other* (type 2 contrast) to capture the same concept of other-regarding (in this case *Both* over *Self*) and non-other regarding (in this case *Bottle* over *Other*) where the chosen options are orthogonal to the preference itself. Notably, these contrasts now instead derive positive ORP from the *Self*/*Both* context (rather than from the *Other*/*Bottle* context) and negative ORP from the *Other*/*Bottle* context (rather than from the *Self*/*Both* context). Spike-field coherence in the *Both*–*Self* contrast exhibited a similar increase in coherence patterns in the beta and gamma band as in the *Other*–*Bottle* contrast (both p > 0.4, Wilcoxon sign rank) (both contrasts examining the effect of delivering juice reward to the conspecific). Similarly, we observed a consistent decrease in the beta and gamma bands both when the actors chose *Self* over *Both* and *Bottle* over *Other* (both p > 0.4) (both contrasts now examining the effect of not delivering juice reward to the conspecific) (**Fig. S7**).

The field-field coherence values for the *Both*–*Self* contrast exhibited a similar increase in their coherence patterns to the beta band as in the *Other*–*Bottle* contrast (both p > 0.22, Wilcoxon sign rank). For the gamma band field-field coherence, however, the *Both*–*Self* contrast showed a more decrease compared to the original contrast (p < 0.0001). We observed a consistent decrease in the beta band field-field coherence both when the actors chose *Self* over *Both* and *Bottle* over *Other* (both p > 0.22, Wilcoxon sign rank), but again showed a more decrease in the gamma band field-field coherence values for *Both*– *Self* contrast compared to the original contrast (p < 0.0001) (**Fig. S8**). These results suggest that the coherence signatures reported were not the mere product of a preferred choice or being in a specific context but were instead driven by the reward outcome of the conspecific monkey.

### Additional Spike-field coherence analyses

To test if the changes in spike-field coherence patterns were not related to simpler sensory-evoked responses, we calculated spike-field coherence values upon the onset of a gray fixation square in the task (from 50 to 150 ms relative to the stimulus onset). We did not observe any significant differences between ACCg_spike_-BLA_field_ and BLA_spike_-ACCg_field_ coherences in both the beta (p = 0.92, Wilcoxon rank sum) and the gamma band (p = 0.36) (**Fig. S9**).

We also performed an additional analysis for the observed spike-field coherence, in which, for each of 1000 iterations, 75% of randomly selected trials were used to recalculate spike-field coherence. These resampled datasets produced consistent results, confirming that our results were not driven by outlier cells, sites, or trials (positive versus negative ORPs: BLA_spike_-ACCg_field_ coherence in the beta band, p = 0.005; ACCg_spike_-BLA_field_ coherence in the gamma band, p = 0.001; Wilcoxon sign rank).

## Supplementary Figures and Legends

**Fig. S1.**
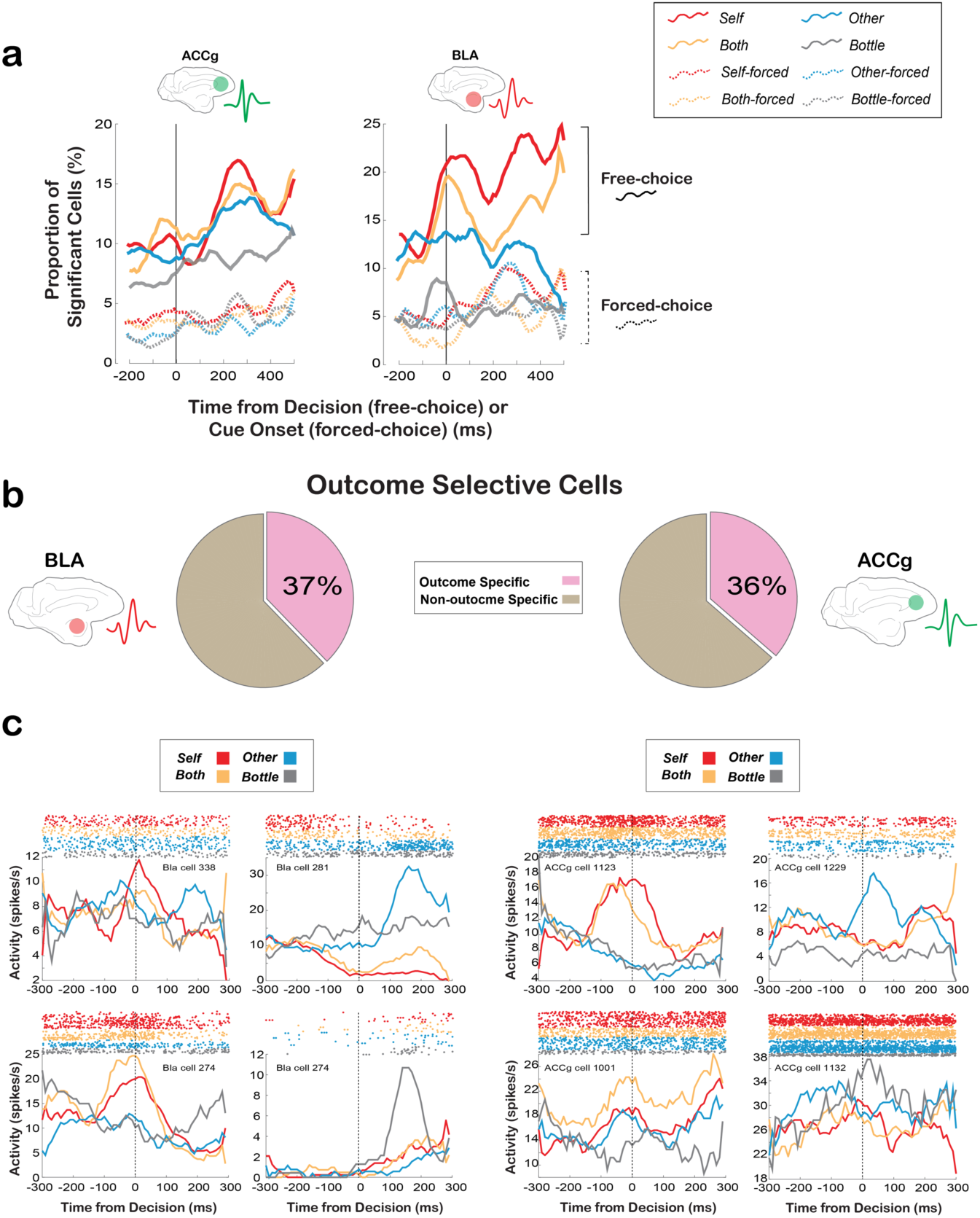
Single neuron summary for encoding *Self*, *Other*, *Both*, and *Bottle* reward outcomes for free-choice and forced-choice trials in BLA and ACCg. (**a**) Proportions of all recorded cells in ACCg (left) and BLA (right) that exhibited significant firing rate modulations in *Self* (purple), *Both* (orange), *Other* (blue), or *Bottle* (gray) condition across time relative to the baseline fixation epoch (p < 0.05, Wilcoxon sign rank). Data from free-choice trials (solid lines) are aligned to the time of decision, whereas data from forced-choice trials (dashed lines) are aligned to the time of cue onset. Note that this analysis examined the time bins in which spikes obtained within *each* outcome condition showed significant difference from its own baseline. (**b**) Pie charts showing the proportions of outcome selective cells in BLA and ACCg. (**c**) Example outcome selective cells in BLA and ACCg. For each area, we show four example cells, each with different outcome-related modulations.

**Fig. S2.**
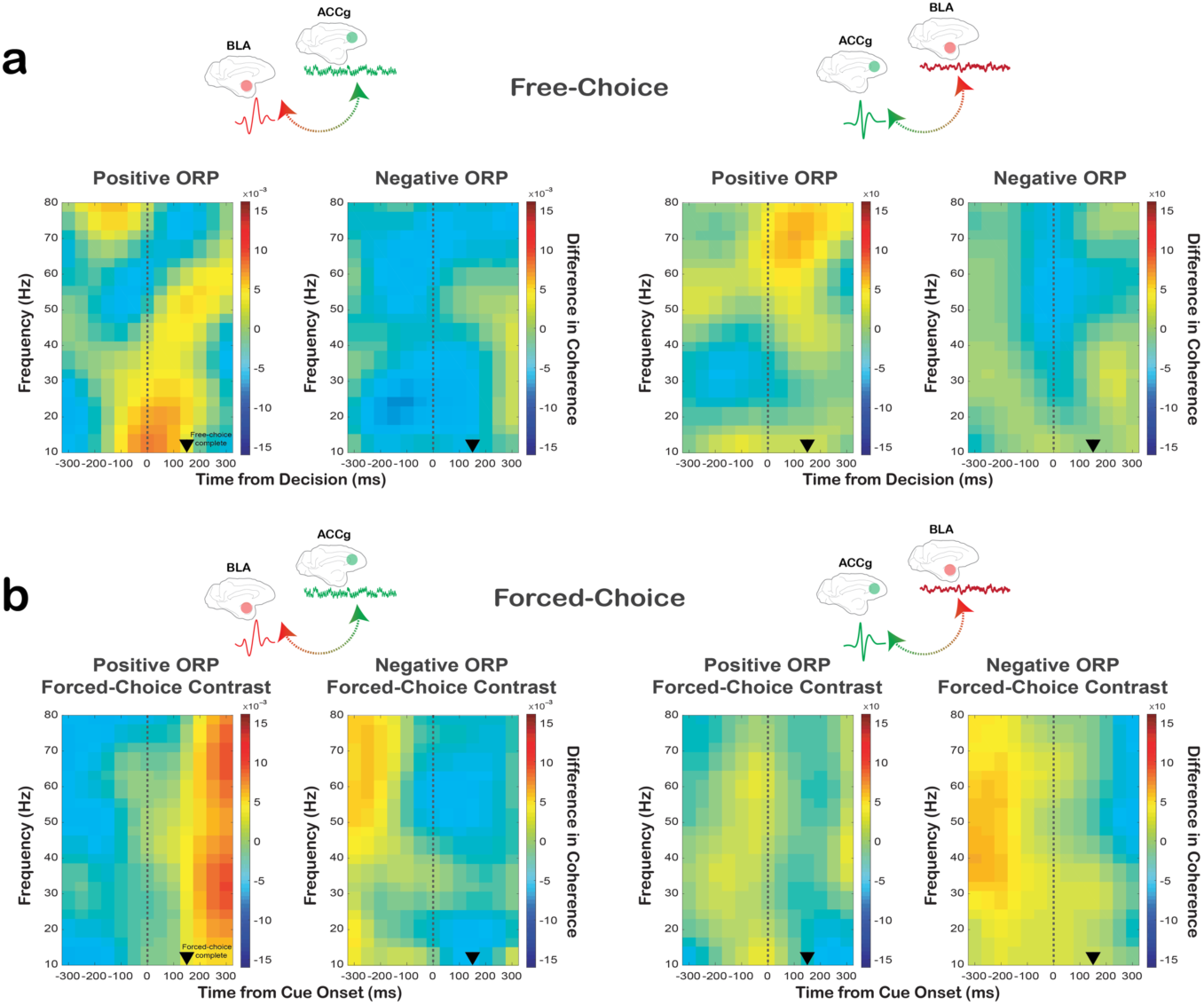
Spike-field coherence between ACCg and BLA cells separately for positive ORP and negative ORP as well as for similar contrasts constructed from forced-choice trials. (**a**) BLA_spike_– ACCg_field_ coherence (left) and ACCg_spike_–BLA_field_ coherence (right) across time and frequency separately for when monkeys actively expressed either positive ORP (choosing *Other* over *Bottle*, *Other*–*Bottle*) or negative ORP (choosing *Self* over *Both*, *Self*–*Both*) on free-choice trials. Data are aligned to the time of decision on free-choice trials. (**b**) BLA_spike_–ACCg_field_ coherence (left) and ACCg_spike_–BLA_field_ coherence (right) across time and frequency separately for when monkeys were presented with computer-determined outcomes on forced-choice trials, contrasting *Other-forced* over *Bottle-forced* to generate positive ORP forced-choice contrast (*Other-forced* – *Bottle-forced*) and *Self-forced* over *Both-forced* to generate negative ORP forced-choice contrast (*Self-forced* – *Both-forced*). Thus, these forced-choice contrasts matched the contrasts used for positive ORP (choosing *Other* over *Bottle*) and negative ORP (choosing *Self* over *Both*) with respect to reward outcome. Data are aligned to the time of cue onset on forced-choice trials. In all plots, the black arrowheads mark the time at which the monkeys completed a free-choice or forced-choice decision by maintaining fixation on a chosen target or cue for 150 ms.

**Fig. S3.**
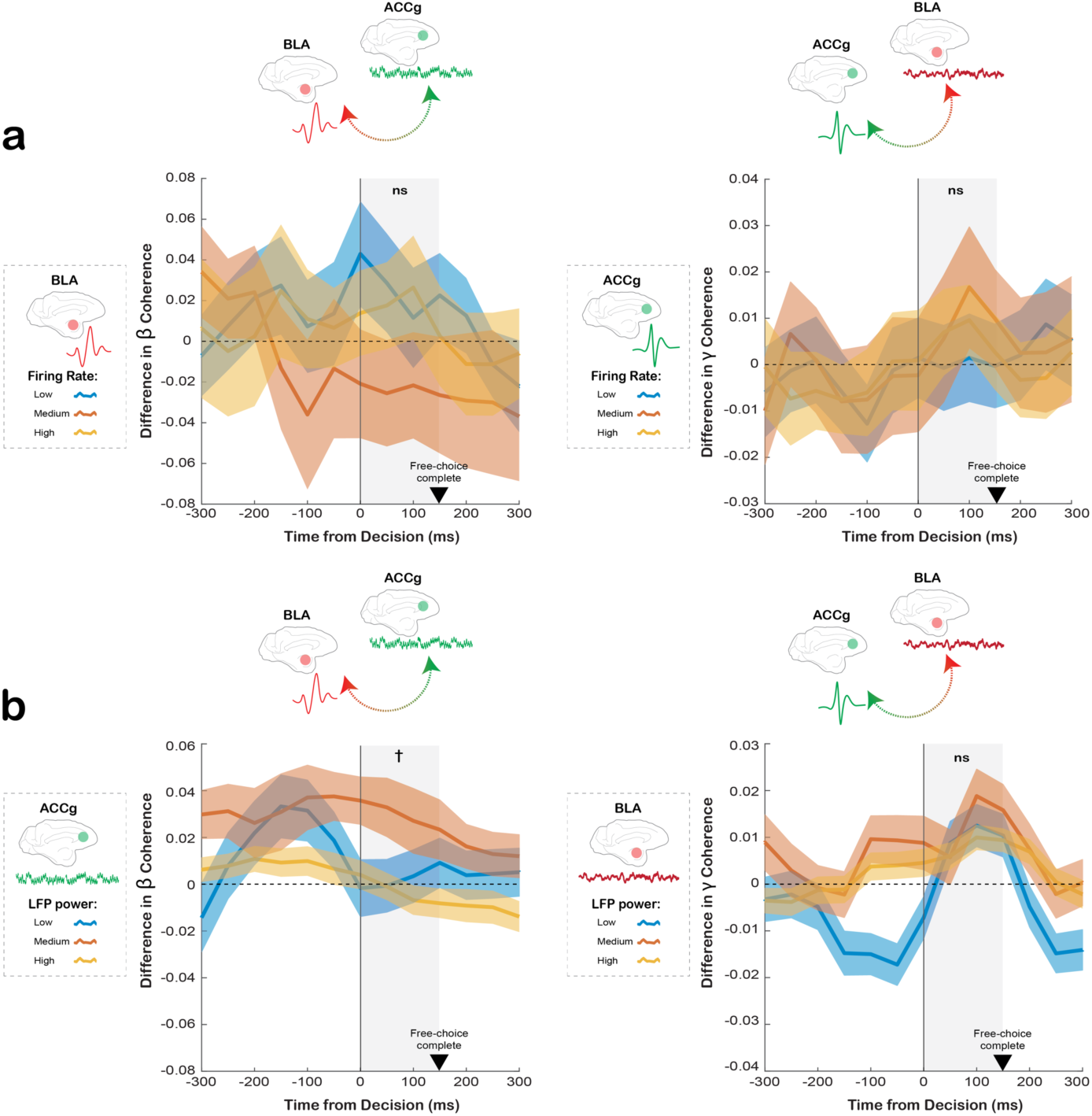
Absence of correlations between the observed spike-field coherence values and the magnitudes of firing rates and LFP power in both BLA and ACCg. (**a**) Differences in the beta BLA_spike_-ACCg_field_ (left) and the gamma ACCg_spike_-BLA_field_ (right) coherence values between positive ORP and negative ORP over time as a function of high, medium, low magnitude quantiles from the firing rates of BLA cells (left) and ACCg cells (right) used in the corresponding spike-field calculations. There were no correlations between the spike-field coherence values across different firing rates of the cells for both comparisons (both p > 0.43, Pearson’s correlation, indicated by ns). (**b**) Differences in the beta BLA_spike_-ACCg_field_ (left) and the gamma ACCg_spike_-BLA_field_ (right) coherence values between positive ORP and negative ORP over time as a function of high, medium, low magnitude quantiles from the LFP powers in ACCg sites (left) and BLA sites (right) used in the corresponding spike-field calculations. There was a weak correlation in the beta band (p = 0.05, indicated by †) but not in the gamma band (p > 0.23, indicated by ns).

**Fig. S4.**
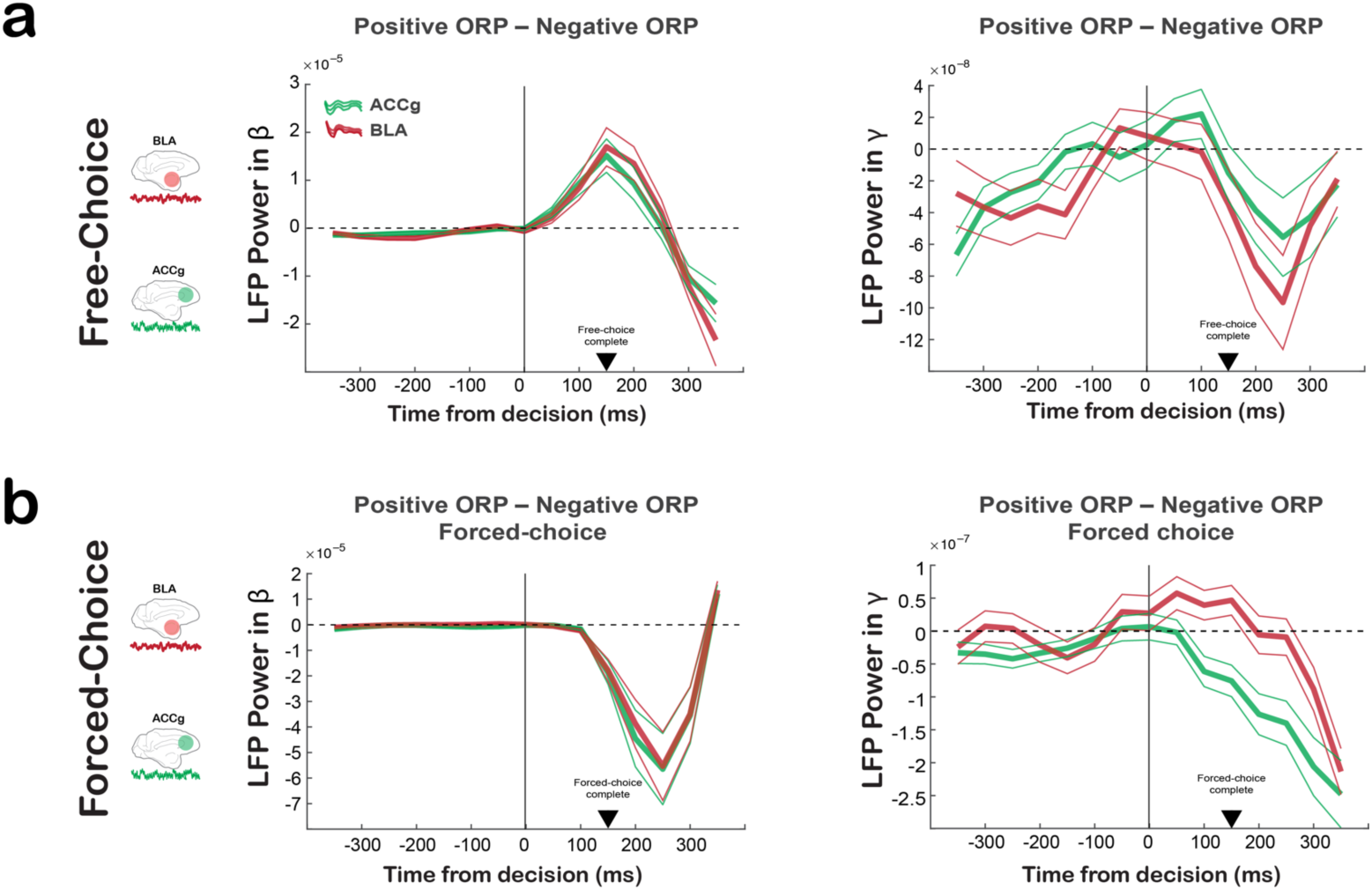
Time courses of LFP powers in the beta and gamma bands in BLA and ACCg with respect to the differences between positive and negative ORPs for both free-choice and forced-choice trials. (**a**) Differences in the beta (left) and the gamma LFP powers (right) between positive ORP and negative ORP over time in BLA and ACCg on free-choice trials. (**b**) Differences in the beta (left) and the gamma LFP powers (right) over time in BLA and ACCg on forced-choice trials, contrasting *Other-forced* over *Bottle-forced* to generate positive ORP forced-choice contrast and *Self-forced* over *Both-forced* to generate negative ORP forced-choice contrast (thus, matching the contrasts across free-choice and forced-choice trials with respect to reward outcome). Except for the gamma band LFP on forced-choice trials (right in the panel **b**), the LFP powers between BLA and ACCg exhibited highly similar modulation time courses. Interestingly, whereas the beta band LFP power differences between the two ORPs markedly increased in both BLA and ACCg following the time of decision on free-choice trials, the same beta band LFP power differences markedly decreased similarly in both areas following cue onset on forced-choice trials.

**Fig. S5.**
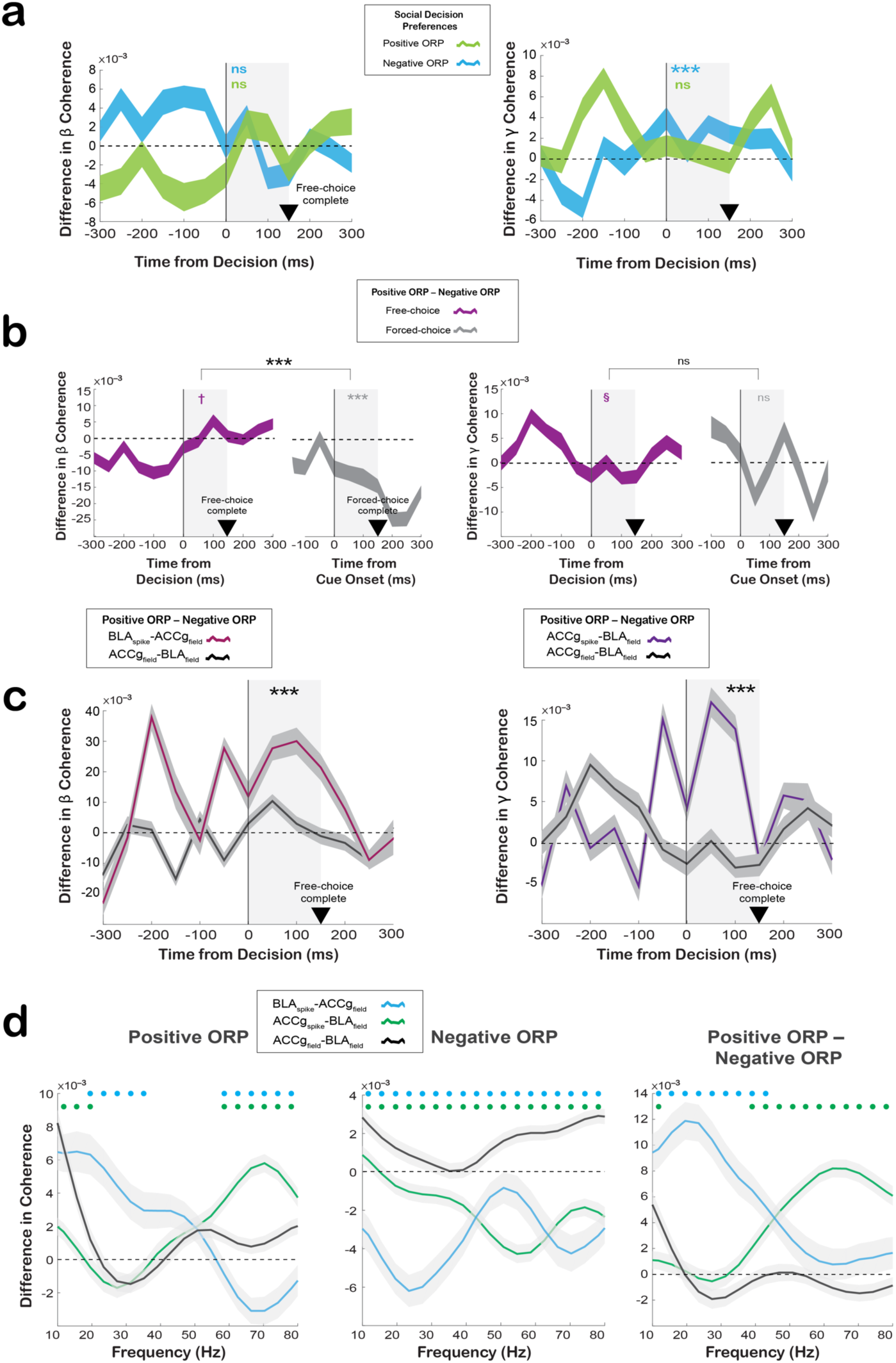
Field-field coherence between ACCg and BLA. (**a**) Time courses of field-field coherence values in the beta frequency (left, 15–25 Hz) and the gamma frequency (right, 45–70 Hz) separately for positive ORP (light green; *Other*–*Bottle*) and negative ORP (light blue; *Self*–*Both*). (**b**) Time courses of beta (left) and gamma (right) band field-field coherence contrasting between the two ORPs on free-choice trials (purple) and forced-choice (forced-choice constructs; see the figure legend in S4) (grey). (**c**) Time courses of coherence differences between the two ORPs on free-choice trials in the beta frequency (left) and the gamma frequency (right) separately for the field-field (dark gray), BLA_spike_-ACC_field_ (dark pink), and ACCg_spike_-BLA_field_ relations (purple). In **a–c**, significant coherence differences between the traces (Wilcoxon rank sum) are indicated in black asterisks for the analyzed epoch (gray shading) (***, p < 0.0001; ns, not significant). In **a–c**, the black arrowheads mark the time at which the monkeys completed a free-choice or forced-choice decision by maintaining fixation on a chosen target or cue for 150 ms. (**d**) Differences between BLA_spike_-ACC_field_ (light blue), ACCg_spike_-ACCg_field_ (green) and ACC_field_-BLA_field_ (dark gray) coherence values across frequency during the post-decision epoch for positive ORP, negative ORP, and for the contrast between the two ORPs. Circles above the lines (in matching colors) show significant differences from the spike-field pairs from the ACC_field_-BLA_field_ coherence values (p < 0.05, Wilcoxon sign rank).

**Fig. S6.**
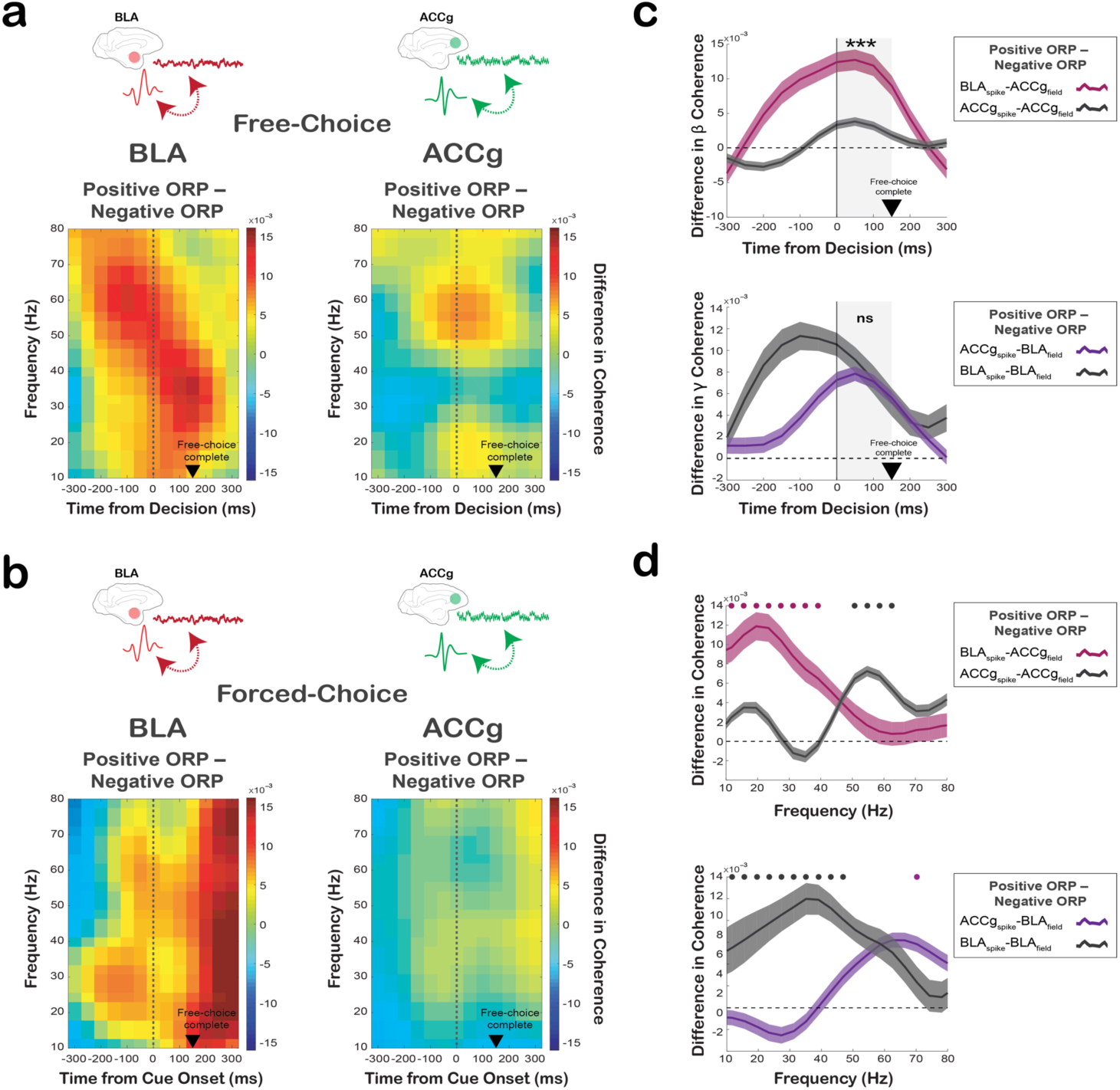
Local, within-region spike-field coherence in ACCg and BLA for positive ORP compared to negative ORP. (**a**) Differences in BLA_spike_-BLA_field_ coherence (left) and ACCg_spike_-ACCg_field_ coherence (right) across time and frequency when monkeys actively expressed positive ORP (choosing *Other* over *Bottle*) versus negative ORP (choosing *Self* over *Both*) on free-choice trials. Data are aligned to the time of free-choice decision. (**b**) Differences in BLA_spike_-BLA_field_ coherence (left) and ACCg_spike_-ACCg_field_ coherence (right) across time and frequency when monkeys were presented with computer-determined outcomes on forced-choice trials, contrasting *Other-forced* over *Bottle-forced* to generate positive ORP forced-choice contrast and *Self-forced* over *Both-forced* to generate negative ORP forced-choice contrast (thus, matching the contrasts across free-choice and forced-choice trials with respect to reward outcome). Data are aligned to the time of cue onset on forced-choice trials. (**c**) (Top) Time courses of spike-field coherence differences between the two ORPs on free-choice trials in the beta band separately for BLA_spike_-ACC_field_ (dark pink) and BLA_spike_-BLA_field_ (dark gray). (Bottom) Time courses of spike-field coherence differences between the two ORPs on free-choice trials in the gamma band separately for ACCg_spike_-BLA_field_ (purple) and ACCg_spike_-ACCg_field_ (dark gray). Significant coherence differences between traces are indicated in black asterisks for the analyzed epoch (gray shading) (***, p < 0.0001; ns, not significant, Wilcoxon rank sum). (**d**) Differences between BLA_spike_-ACC_field_ (dark pink) and BLA_spike_-BLA_field_ (dark gray) coherence values across frequency during the post-decision epoch (top) and between ACCg_spike_-BLA_field_ (purple) and ACCg_spike_-ACCg_field_ (dark gray) values across frequency during the post-decision epoch (bottom). Circles above the lines (in matching colors) show significant differences from the spike-field pairs from the ACCg_field_-BLA_field_ coherence values (p < 0.05, Wilcoxon sign rank). In **a**–**c**, the black arrowheads mark the time at which the monkeys completed a free-choice or a forced-choice decision by maintaining the fixation on a target or cue for 150 ms.

**Fig. S7.**
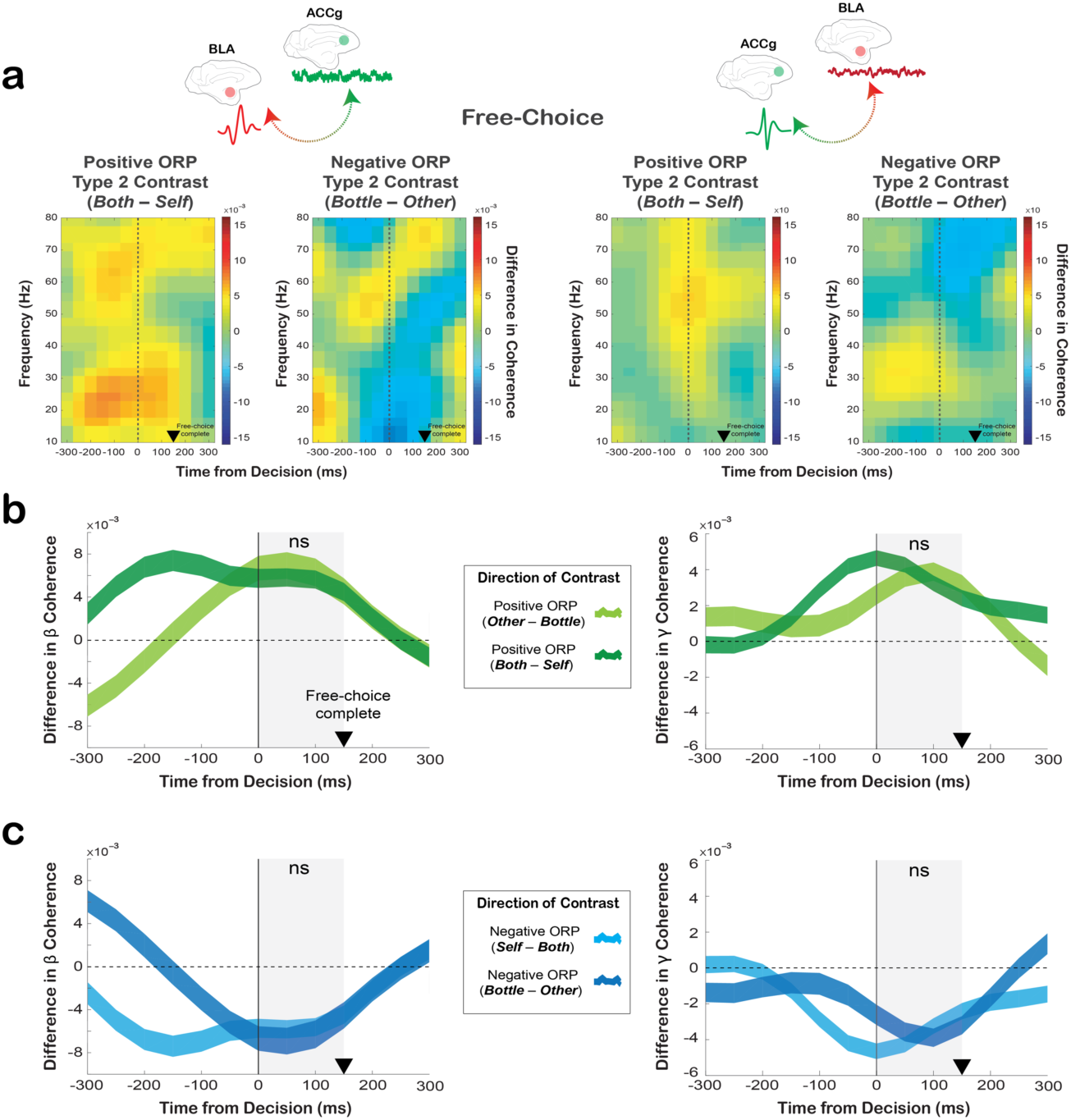
Different ways of contrasting positive and negative ORPs (type 2 contrasts) replicate the main spike-field coherence findings. (**a**) Free-choice spectograms showing the BLA_spike_-ACCg_field_ coherence (left two panels) and ACCg_spike_-BLA_field_ coherence (right two panels) values after applying the type 2 contrasts (positive ORP now derived from the *Self*/*Both* context as *Both*–*Self*; negative ORP now derived from the *Other*/*Bottle* context as *Bottle*–*Other*). (**b**) Comparisons over time for the beta BLA_spike_-ACCg_field_ coherence (left) and the gamma ACCg_spike_-BLA_field_ coherence (right) values between the original positive ORP contrast (*Other*–*Bottle*) and the type 2 positive ORP contrast (*Both*–*Self*). Although there were differences between the two contrasts prior to making a choice, the two contrasts yielded comparable coherence values in both comparisons (both p > 0.41, Wilcoxon sign rank). (**c**) Comparisons over time for the beta BLA_spike_-ACCg_field_ coherence (left) and the gamma ACCg_spike_-BLA_field_ coherence (right) values between the original negative ORP contrast (*Self*–*Both*) and the type 2 negative ORP contrast (*Bottle*–*Other*). Although there were differences between the two contrasts prior to making a choice, the two contrasts yielded comparable coherence values in both comparisons (both p > 0.41).

**Fig. S8.**
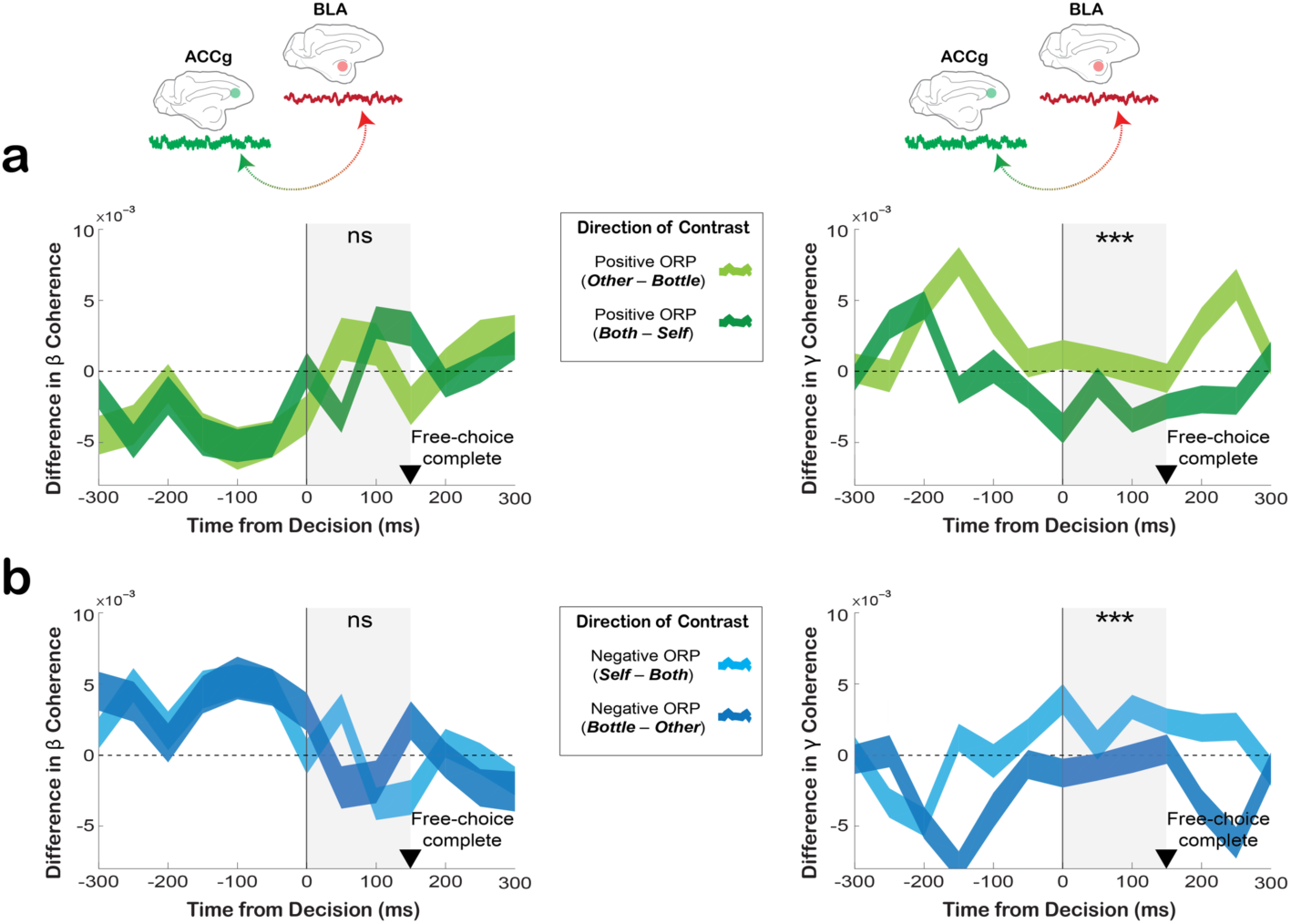
Different ways of contrasting positive and negative ORPs (type 2 contrasts) for the field-field coherence between BLA and ACCg. (**a**) Comparisons over time for the beta BLAspike-ACCgfield coherence (left) and the gamma BLA_field_-ACCg_field_ coherence (right) values between the original positive ORP contrast (*Other*–*Bottle*) and the type 2 positive ORP contrast derived from the *Self*/*Both* context (*Both*–*Self*). No coherence differences were found in the beta band (p > 0.22, Wilcoxon sign rank). In the gamma band, the type 2 contrast showed reduced coherence values in the main analysis epoch (grey shading) compared to the original contrast (p < 0.0001). (**b**) Comparisons over time for the beta BLA_field_-ACCg_field_ coherence (left) and the gamma BLA_field_-ACCg_field_ coherence (right) values between the original negative ORP contrast (*Self*–*Both*) and the type 2 negative ORP contrast derived from the *Other*/*Both* context (*Bottle*–*Other*). Again, no coherence differences were found in the beta band (p > 0.22), but the original contrast showed increased coherence values in the main analysis epoch (grey shading) compared to the type 2 contrast (p < 0.0001).

**Fig. S9.**
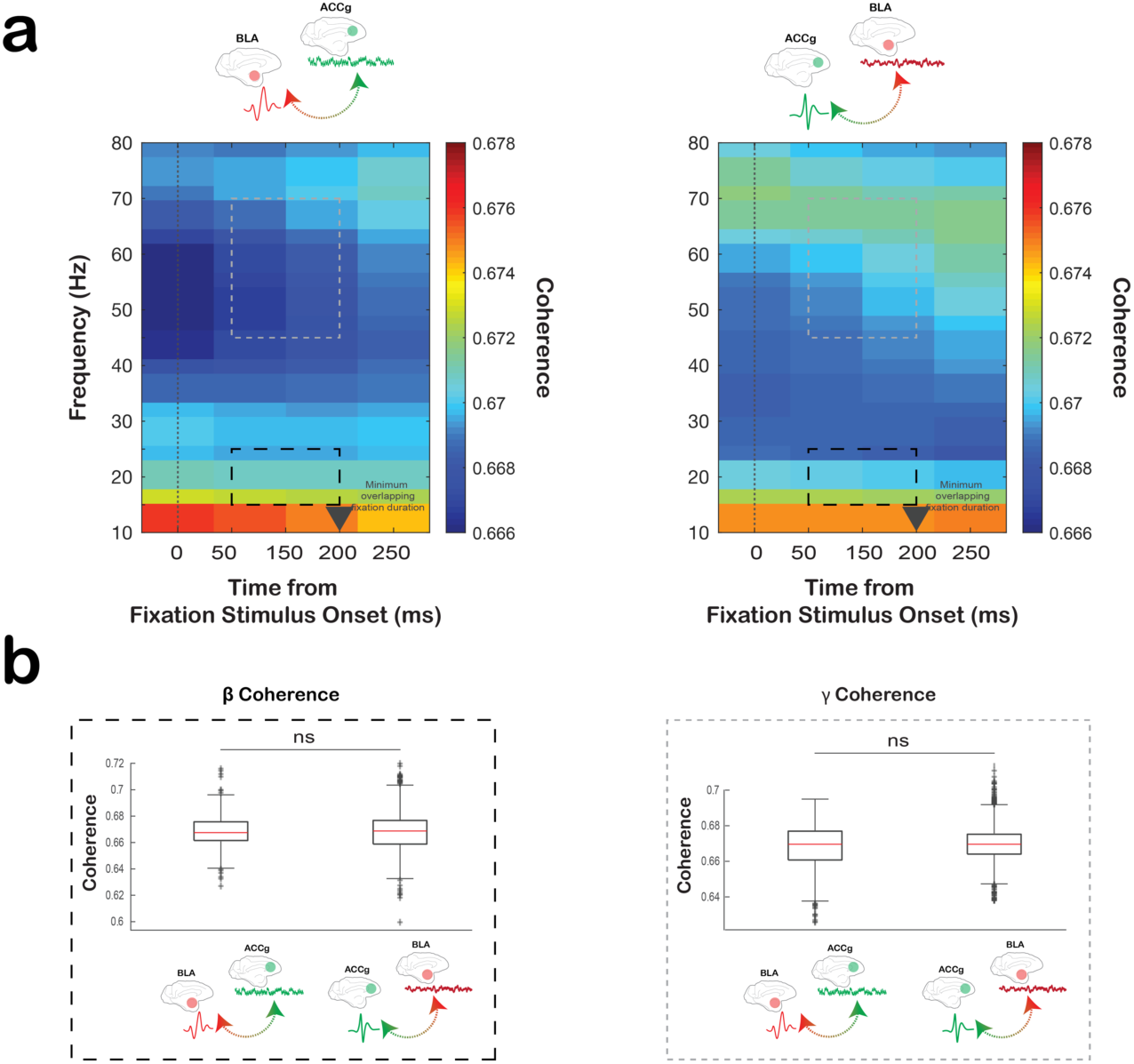
Spike-field coherence values between BLA and ACCg upon the onset of visual fixation stimulus. (**a**) Spectograms showing the BLA_spike_-ACCg_field_ coherence (left) and ACCg_spike_-BLA_field_coherence (right) values aligned to the onset of fixation stimulus (grey fixation square). Monkeys were required to fixate on this stimulus upon its onset for at least 200 ms. (**b**) Quantifications of the beta (left, 15–25Hz) and gamma (right, 45–75Hz) coherence values in the 50–200 ms period relative to the fixation stimulus onset between the BLA_spike_-ACCg_field_ coherence and ACCg_spike_-BLA_field_ coherence values. No differential coherence values were observed between the two coherence pairs for both frequency bands (beta p > 0.92, gamma p > 0.36, Wilcoxon rank sum).

**Fig. S10.**
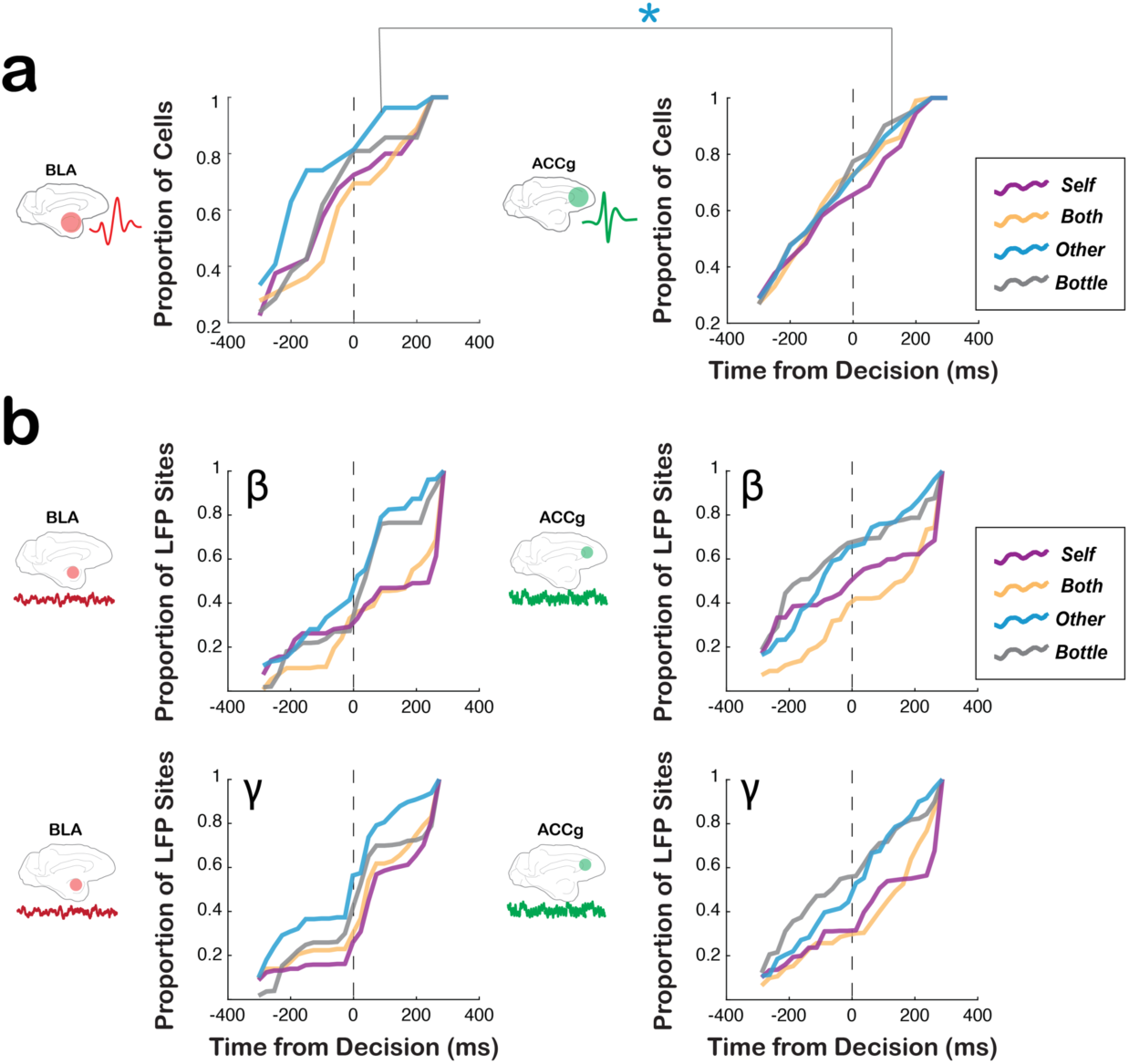
Emergence times for outcome-related signals for spikes and LFP powers in BLA and ACCg. (**a**) Cumulative histograms showing the time points at which BLA cells (left) and ACCg cells (right) began to show significant outcome-related signals, separately for *Self*, *Both*, *Other*, and *Bottle*. Between BLA and ACCg, only *Other* choice signals emerged earlier in BLA than ACCg (*, p < 0.001, Kolmogorov–Smirnov test). All the other comparisons between the two areas had comparable cumulative distributions (all, p > 0.08). (**b**) Cumulative histograms showing the time points at which BLA (left) and ACCg LFP sites (right) began to show significant outcome-related LFP power signals compared to baseline, separately for *Self*, *Both*, *Other*, and *Bottle*, and separately for the beta (top row) and gamma bands (bottom row). All comparisons within the same frequency band between the two areas had comparable cumulative distributions (all, p > 0.38).

**Fig. S11.**
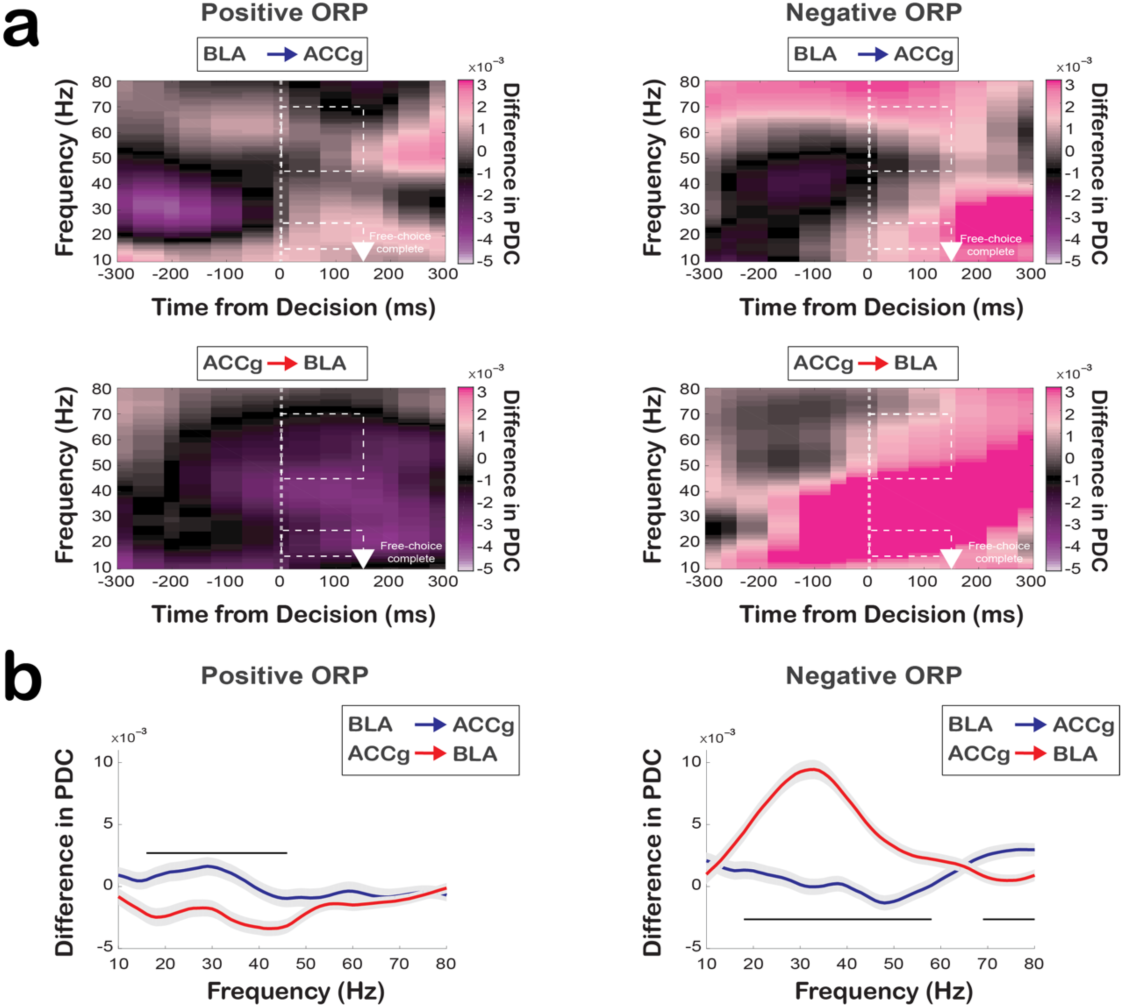
Directionality of information flow between BLA and ACCg for computer-determined outcomes on forced-choice trials as a function of time and frequency. (**a**) Frequency-domain directional influences assessed by partial directed coherence (PDC) when monkeys were presented with computer-determined outcomes (forced-choice trials). PDC values as a function of time and frequency for *Other-forced* over *Bottle-forced* (positive ORP forced-choice contrast; see the legend in **Fig. S2**) for BLA➔ACCg (top left) and ACCg➔BLA (bottom left), and PDC values for *Self-forced* over *Both-forced* (negative ORP forced-choice contrast) for BLA➔ACCg (top right) and ACCg➔BLA (bottom right). Data are aligned to the time of cue onset. The white arrowheads mark the time at which the monkeys completed a forced-choice decision by maintaining fixation on a chosen target for 150 ms. Dotted lines indicate the beta (15–25Hz) and gamma (45–70Hz) band for the post-decision epoch. (**b**) Quantifications of the directional information flow on forced-choice trials as a function of frequency for positive ORP decision (left) and negative ORP (right) for BLA➔ACCg (in blue) and ACCg➔BLA (in red). In b horizontal lines indicate significant different between BLA➔ACCg and ACCg➔BLA (p < 0.05, Wilcoxon sign rank). Shaded regions represent standard errors.

## Notes

#### Summary of Updates

This is a revised manuscript.

## References

1. Behrens, T. E. J., Hunt, L. T. & Rushworth, M. F. S. The computation of social behavior. Science 324, 1160–1164 (2009).

2. Bhanji, J. P. & Delgado, M. R. The social brain and reward: social information processing in the human striatum. Wiley Interdiscip. Rev. Cogn. Sci. 5, 61–73 (2014).

3. Sliwa, J. & Freiwald, W. A. A dedicated network for social interaction processing in the primate brain. Science 356, 745–749 (2017).

4. Ruff, C. C. & Fehr, E. The neurobiology of rewards and values in social decision making. Nat. Rev. Neurosci. 15, 549–562 (2014).

5. Seo, H. & Lee, D. Neural basis of learning and preference during social decision-making. Curr. Opin. Neurobiol. 22, 990–995 (2012).

6. Chang, S. W. C., Gariépy, J.-F. & Platt, M. L. Neuronal reference frames for social decisions in primate frontal cortex. Nat. Neurosci. 16, 243–250 (2013).

7. Haroush, K. & Williams, Z. M. Neuronal prediction of opponent’s behavior during cooperative social interchange in primates. Cell 160, 1233–1245 (2015).

8. Noritake, A., Ninomiya, T. & Isoda, M. Social reward monitoring and valuation in the macaque brain. Nat. Neurosci. 21, 1452–1462 (2018).

9. Chang, S. W. C. et al. Neural mechanisms of social decision-making in the primate amygdala. Proc. Natl. Acad. Sci. 112, 16012–16017 (2015).

10. Grabenhorst, F., Báez-Mendoza, R., Genest, W., Deco, G. & Schultz, W. Primate Amygdala neurons simulate decision processes of social partners. Cell 177, 986–998.e15 (2019).

11. Munuera, J., Rigotti, M. & Salzman, C. D. Shared neural coding for social hierarchy and reward value in primate amygdala. Nat. Neurosci. 21, 415–423 (2018).

12. Azzi, J. C. B., Sirigu, A. & Duhamel, J.-R. Modulation of value representation by social context in the primate orbitofrontal cortex. Proc. Natl. Acad. Sci. 109, 2126–2131 (2012).

13. Baez-Mendoza, R., Harris, C. J. & Schultz, W. Activity of striatal neurons reflects social action and own reward. Proc. Natl. Acad. Sci. 110, 16634–16639 (2013).

14. Falcone, R., Brunamonti, E., Ferraina, S. & Genovesio, A. Neural encoding of self and another agent’s goal in the primate prefrontal cortex: human-monkey interactions. Cereb. Cortex 26, 4613– 4622 (2016).

15. Nummela, S. U., Jovanovic, V., Mothe, L. de la & Miller, C. T. Social Context-dependent activity in marmoset frontal cortex populations during natural conversations. J. Neurosci. 37, 7036–7047 (2017).

16. Apps, M. A. J., Rushworth, M. F. S. & Chang, S. W. C. The anterior cingulate gyrus and social cognition: tracking the motivation of others. Neuron 90, 692–707 (2016).

17. Hill, M. R., Boorman, E. D. & Fried, I. Observational learning computations in neurons of the human anterior cingulate cortex. Nat. Commun. 7, 12722 (2016).

18. Zaki, J. & Ochsner, K. The neuroscience of empathy: progress, pitfalls and promise. Nat. Neurosci. 15, 675–680 (2012).

19. Mars, R. B. et al. On the relationship between the “default mode network” and the “social brain”. Front. Hum. Neurosci. 6, (2012).

20. Amadei, E. A. et al. Dynamic corticostriatal activity biases social bonding in monogamous female prairie voles. Nature 546, 297–301 (2017).

21. Allsop, S. A. et al. Corticoamygdala transfer of socially derived information gates observational learning. Cell 173, 1329–1342.e18 (2018).

22. Zhan, Y. et al. Deficient neuron-microglia signaling results in impaired functional brain connectivity and social behavior. Nat. Neurosci. 17, 400–406 (2014).

23. Carmichael, S. T. & Price, J. L. Limbic connections of the orbital and medial prefrontal cortex in macaque monkeys. J. Comp. Neurol. 363, 615–641 (1995).

24. Klavir, O., Genud-Gabai, R. & Paz, R. Functional connectivity between amygdala and cingulate cortex for adaptive aversive learning. Neuron 80, 1290–1300 (2013).

25. Pesaran, B. et al. Investigating large-scale brain dynamics using field potential recordings: analysis and interpretation. Nat. Neurosci. 21, 903 (2018).

26. Fries, P. A mechanism for cognitive dynamics: neuronal communication through neuronal coherence. Trends Cogn. Sci. 9, 474–480 (2005).

27. Chang, S. W. C., Winecoff, A. A. & Platt, M. L. Vicarious reinforcement in rhesus macaques (*Macaca mulatta*). Front. Neurosci. 5, (2011).

28. Chang, S. W. C., Barter, J. W., Ebitz, R. B., Watson, K. K. & Platt, M. L. Inhaled oxytocin amplifies both vicarious reinforcement and self reinforcement in rhesus macaques (*Macaca mulatta*). Proc. Natl. Acad. Sci. 109, 959–964 (2012).

29. Paxinos, G., Huang, X.-F. & Toga, A. W. The Rhesus Monkey Brain in Stereotaxic Coordinates. (Academic Press, 1999).

30. Baccalá, L. A. & Sameshima, K. Partial directed coherence: a new concept in neural structure determination. Biol. Cybern. 84, 463–474 (2001).

31. Buzsáki, G. & Wang, X.-J. Mechanisms of gamma oscillations. Annu. Rev. Neurosci. 35, 203–225 (2012).

32. Fries, P. Rhythms for cognition: Communication through coherence. Neuron 88, 220–235 (2015).

33. Pesaran, B. et al. Investigating large-scale brain dynamics using field potential recordings: analysis and interpretation. Nat. Neurosci. 21, 903–919 (2018).

34. Hipp, J. F., Engel, A. K. & Siegel, M. Oscillatory synchronization in large-scale cortical networks predicts perception. Neuron 69, 387–396 (2011).

35. Womelsdorf, T., Fries, P., Mitra, P. P. & Desimone, R. Gamma-band synchronization in visual cortex predicts speed of change detection. Nature 439, 733–736 (2006).

36. Wong, Y. T., Fabiszak, M. M., Novikov, Y., Daw, N. D. & Pesaran, B. Coherent neuronal ensembles are rapidly recruited when making a look-reach decision. Nat. Neurosci. 19, 327–334 (2016).

37. Kahana, M. J., Sekuler, R., Caplan, J. B., Kirschen, M. & Madsen, J. R. Human theta oscillations exhibit task dependence during virtual maze navigation. Nature 399, 781–784 (1999).

38. Fujisawa, S. & Buzsáki, G. A 4 Hz oscillation adaptively synchronizes prefrontal, VTA, and hippocampal activities. Neuron 72, 153–165 (2011).

39. Adhikari, A., Topiwala, M. A. & Gordon, J. A. Synchronized activity between the ventral hippocampus and the medial prefrontal cortex during anxiety. Neuron 65, 257 (2010).

40. Antzoulatos, E. G. & Miller, E. K. Increases in functional connectivity between prefrontal cortex and striatum during category learning. Neuron 83, 216–225 (2014).

41. Brincat, S. L. & Miller, E. K. Frequency-specific hippocampal-prefrontal interactions during associative learning. Nat. Neurosci. 18, 576–581 (2015).

42. Taub, A. H., Perets, R., Kahana, E. & Paz, R. Oscillations synchronize amygdala-to-prefrontal primate circuits during aversive learning. Neuron 97, 291–298.e3 (2018).

43. Buschman, T. J. & Miller, E. K. Top-down versus bottom-up control of attention in the prefrontal and posterior parietal cortices. Science 315, 1860–1862 (2007).

44. Engel, A. K., Fries, P. & Singer, W. Dynamic predictions: oscillations and synchrony in top-down processing. Nat. Rev. Neurosci. 2, 704–716 (2001).

45. Engel, A. K. & Fries, P. Beta-band oscillations--signalling the status quo? Curr. Opin. Neurobiol. 20, 156–165 (2010).

46. Cardin, J. A. et al. Driving fast-spiking cells induces gamma rhythm and controls sensory responses. Nature 459, 663–667 (2009).

47. Jia, X. & Kohn, A. Gamma rhythms in the brain. PLOS Biol. 9, e1001045 (2011).

48. Livneh, U., Resnik, J., Shohat, Y. & Paz, R. Self-monitoring of social facial expressions in the primate amygdala and cingulate cortex. Proc. Natl. Acad. Sci. 109, 18956–18961 (2012).

49. Gothard, K. M., Battaglia, F. P., Erickson, C. A., Spitler, K. M. & Amaral, D. G. Neural responses to facial expression and face identity in the monkey amygdala. J. Neurophysiol. 97, 1671–1683 (2007).

50. Grabenhorst, F., Hernádi, I. & Schultz, W. Prediction of economic choice by primate amygdala neurons. Proc. Natl. Acad. Sci. 109, 18950–18955 (2012).

51. Liu, Y., Yttri, E. A. & Snyder, L. H. Intention and attention: different functional roles for LIPd and LIPv. Nat. Neurosci. 13, 495–500 (2010).

52. Chang, S. W. et al. Neural mechanisms of social decision-making in the primate amygdala. Proc. Natl. Acad. Sci. 112, 16012–16017 (2015).

53. Chung, J. E. et al. A fully automated approach to spike sorting. Neuron 95, 1381–1394.e6 (2017).

54. Bokil, H., Andrews, P., Kulkarni, J. E., Mehta, S. & Mitra, P. P. Chronux: a platform for analyzing neural signals. J. Neurosci. Methods 192, 146–151 (2010).

55. Jarvis, M. R. & Mitra, P. P. Sampling properties of the spectrum and coherency of sequences of action potentials. Neural Comput. 13, 717–749 (2001).

56. Sommerlade, L. et al. Time-variant estimation of directed influences during Parkinsonian tremor. J. Physiol. Paris 103, 348–352 (2009).

57. Omidvarnia, A., Azemi, G., Boashash, B., O’Toole, J. M., Colditz, P. B., & Vanhatalo, S. (2013). Measuring time-varying information flow in scalp EEG signals: orthogonalized partial directed coherence. IEEE transactions on biomedical engineering, 61(3), 680–693.

58. Saez, A., Rigotti, M., Ostojic, S., Fusi, S. & Salzman, C. D. Abstract Context Representations in Primate Amygdala and Prefrontal Cortex. Neuron 87, 869–881 (2015).

59. Sutton, R. S., & Barto, A. G. Reinforcement learning: An introduction. MIT press. (2018)

